# Lysine-deficient proteome can be regulated through non-canonical ubiquitination and ubiquitin-independent proteasomal degradation

**DOI:** 10.1101/2023.01.18.524605

**Authors:** Natalia A. Szulc, Małgorzata Piechota, Pankaj Thapa, Wojciech Pokrzywa

## Abstract

The ubiquitin-proteasome system (UPS) removes damaged and unwanted proteins by attaching ubiquitin to lysines in a process termed ubiquitination. Little is known how functional components of the UPS, often exposed to erroneous labeling by ubiquitin during functioning, avoid premature proteolysis. An extensive lysine-less region (lysine desert) in the yeast E3 ligase Slx5 was shown to counteract its ubiquitin-dependent turnover. We conducted bioinformatic screens among prokaryotes and eukaryotes to describe the scope and conservation of this phenomenon. We found that lysine deserts are widespread among bacteria using pupylation-dependent proteasomal degradation, an analog of the UPS. In eukaryotes, lysine deserts appear with increasing organismal complexity, and the most evolutionarily conserved are enriched in the UPS members. Using VHL and SOCS1 E3 ligases, which elongate their lysine desert in the course of evolution, we established that they are non-lysine ubiquitinated, which does not influence their stability, and can be subject to proteasome turnover irrespective of ubiquitination. Our data suggest that a combination of non-lysine ubiquitination and ubiquitin-independent degradation may control the function and fate of the lysine-deficient proteome, as the presence of lysine deserts does not correlate with the half-life.

## Introduction

Maintaining protein homeostasis (proteostasis) requires the degradation of damaged or unwanted proteins and plays a crucial role in cellular function, organismal growth, and, ultimately, viability^1, 2^. A principal proteolytic component of the cellular proteostasis network is the ubiquitin-proteasome system (UPS)^3^. Enzymes operating within the UPS recognize the substrates destined for degradation and label them by covalent attachment of a small, evolutionarily conserved protein – ubiquitin^4^. Ubiquitination of a substrate requires a cascade of enzymes. The ubiquitin-activating enzyme (E1) hydrolyzes adenosine triphosphate (ATP) and forms a high-energy thioester bond between an internal cysteine residue and the C-terminus of ubiquitin, containing a di-glycine (diGly) motif. Activated ubiquitin is then passed on to the ubiquitin-conjugating enzyme (E2), which forms similar thioester-linked complexes. Finally, ubiquitin is covalently attached mainly to lysine sidechains of the substrate protein by a ubiquitin ligase (E3), which often directly interacts with the substrate^5^.

Mechanistically, two classes of E3s are widely characterized. HECT (Homologous to E6AP C-terminal) E3s form an intermediate thioester bond with the ubiquitin before catalyzing substrate ubiquitination. By contrast, RING (Really Interesting New Gene)/U-box E3s are molecular scaffolds that bring E2-ubiquitin and target protein in close proximity to facilitate the transfer of ubiquitin^6–8^. The majority of E3s belong to the RING family, with the largest subfamily being the cullin-RING ligases (CRLs), which have more than 400 members and are accountable for approximately 20% of the total ubiquitination in cells^9^.

A single ubiquitination event usually does not direct the protein to degradation^10, 11^. Instead, labeling with polyubiquitin chains, formed by an isopeptide linkage between the preceding ubiquitin’s lysine and the next subunit’s C-terminal glycine, acts as a potent signal of protein destruction. Polyubiquitin chains can be homotypic (involving the same lysine residue), heterotypic (involving different lysine residues in one chain), or branched, in which the ubiquitin molecule is subsequently ubiquitinated on different lysine positions^10, 11^. Chains of four to six ubiquitin moieties linked via lysine residues 48 of ubiquitin usually promote degradation of modified substrates by the 26S proteasome - a multi-catalytic protease complex composed of a barrel-shaped 20S proteolytic core particle and a 19S regulatory particle that translocates substrates into the former where they are degraded into short peptides^10–13^. Moreover, ubiquitination can also occur on the free amino group of the N-terminus of a protein, as well as on serine, cysteine, and threonine residues^14, 15^. This non-canonical ubiquitination, although thermodynamically less stable^14, 16, 17^ than the modification on lysine, also targets proteins for proteasomal degradation^16^. However, ubiquitination is not an irreversible modification - deubiquitinating enzymes (DUBs) can cleave ubiquitin from the protein substrate and revert the fate of the ubiquitin-conjugated proteins^18^, edit ubiquitin chains, and process ubiquitin precursors^19^. Recently, it has also been shown that some DUBs have high esterase activity and preferentially cleave cysteine-conjugated ubiquitin^20^, implying the complexity and importance of the unexplored code of non-canonical ubiquitination.

A process analogous to eukaryotic ubiquitination was identified in *Actinobacteria*. Here, the lysine residues of bacterial target proteins are covalently modified with a small protein called Pup (a prokaryotic ubiquitin-like protein) in a process called pupylation^21^. Although Pup and ubiquitin share functional analogies^22^, they are neither homologous in amino acid sequence nor structure^23^. A feature shared with ubiquitin is the conserved diGly motif; however, it is not located at the C-terminus itself, as in processed ubiquitin, but followed by either glutamine or glutamic acid in all Pup sequences. A single enzyme, the Pup ligase, PafA^24, 25^ catalyses Pup attachment to substrates via isopeptide bonding. Pupylation is counterbalanced by a depupylating enzyme, Dop, that mediates isopeptide bond cleavage, releasing Pup from the modified protein^26, 27^. The proteasome complex recognizes pupylated proteins due to the binding of Pup to the N-terminal coiled-coil domains of the proteasomal ATPase^28, 29^. Pupylation-dependent proteasomal degradation is restricted to the *Actinobacteria* phylum, including pathogens of importance such as *M. tuberculosis* and *M. leprae*^21^.

E3s, while explicitly recognizing their target proteins, are known to be promiscuous when choosing the lysine residues and often modify multiple substrate positions in their region of action (ubiquitination zone)^30^. Furthermore, in the absence of target proteins, E3s can self-catalyze ubiquitination (auto-ubiquitination), leading to their degradation^31^. For some, self-initiated degradation is regulated by substrate availability and is probably a desirable feature evolutionarily consolidated^32^. Other E3s avoid this fate by recruiting suitable DUBs; examples include the protection of RNF123 by USP19^33^ or the stabilization of viral ICP0 by recruiting cellular USP7^34^. E3s can also be targeted for degradation by ubiquitination performed by other E3s^31^. However, E3s targeting misfolded proteins do not require such moderation; their auto- ubiquitination and subsequent degradation would deplete the cell of valuable quality control factors^31^. Thus, proteolytic mechanisms may also threaten essential or functional proteins, including proteins that constitute the architecture of the UPS. Therefore, functional proteomes must have evolved mechanisms to prevent unintended proteolytic destruction.

An intuitive “strategy” of a proteome susceptible to premature ubiquitination is to avoid lysines in critical domains or entire sequences, potentially leaving a few whose ubiquitination can be precisely controlled. Indeed, an extensive lysine-less region (lysine desert) in the yeast SUMO-targeted ubiquitin ligase (STUbL) Slx5, constituting over 64% of its sequence, was shown to counteract its ubiquitin-dependent turnover^35^. A similar example is the San1 yeast E3 ligase, in which self-destruction is limited by the lack of lysines in disordered substrate-binding regions that could be immediately ubiquitinated after binding to the E2-ubiquitin complex^36^. Extensive lysine deserts are also found in other protein quality control factors, including BAG6^37^. Moreover, many bacterial AB-type toxins exhibit unusual lysine depletion in their sequence^38^, presumably to avoid ubiquitination and degradation. Thus, lysine desert could be common in various organisms’ proteomes and constitute an adaptation of functional proteomes to avoid premature turnover.

To measure the widespread of lysine desert phenomenon, we conducted bioinformatic screens among prokaryotes and eukaryotes, considering not only mere lysine desert appearance in sequence but also its conservation in orthologs, which allowed us to gain deeper insights into the evolutionary traits of this phenomenon and discover conserved lysine desert regions, pointing at their possible functional role. We also assessed *in cellula* the role of lysine desert in preventing the degradation of VHL (von Hippel-Lindau tumor suppressor) and SOCS1 (suppressor of cytokine signaling 1) substrate receptor subunits of the cullin-RING E3 complexes^39^. We demonstrated that the lysine deserts in VHL and SOCS1 undergo non-canonical ubiquitination, which likely serves regulatory-only purposes. On the other hand, their stability is proteasome-dependent but ubiquitination-independent, which probably accounts for the lack of correlation between lysine desert and the half-life of the equipped human proteome.

## Results

We first aimed to define the lysine desert region regarding the absolute amino acid (aa) length and sequence fraction. To find such thresholds, we searched for continuous lysine-less regions using sliding windows of varying lengths, defined as either sequence fraction (Fig 1A) or nominal value (Fig 1B), in proteins (≥150 aa) from all UniProt^40^ eukaryotic reference proteomes (1329 taxons; Table S1). Based on these results, we defined the lysine desert as a continuous lysine-less region either constituting min. 50% of a given sequence (hereafter lysine desert min. 50%) or min. 150 aa length (hereafter lysine desert min. 150 aa) (Fig 1C), as such defined lysine deserts occur in less than 10% of eukaryotic proteins.

**Figure 1.**
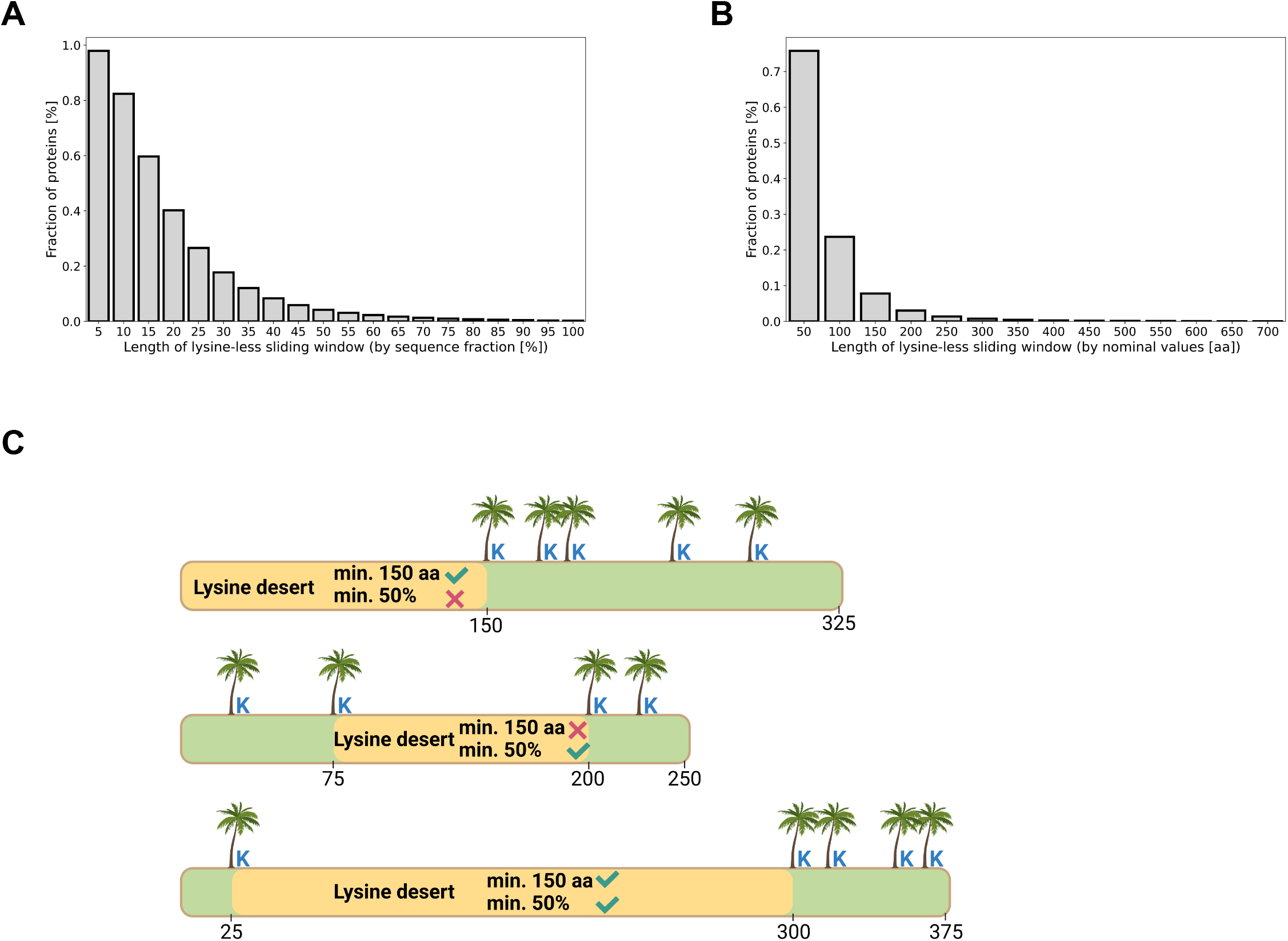
Definition of a lysine desert. **(A)** Fraction of occurrence of lysine-less regions, probed by sliding windows of varying lengths defined as sequence fraction, among sequences (≥150 aa long) from all UniProt eukaryotic reference proteomes. **(B)** Fraction of occurrence of lysine-less regions, probed by sliding windows of varying lengths defined as the multiplicity of 50, among sequences (≥150 aa long) from all UniProt eukaryotic reference proteomes. **(C)** Two definitions of a lysine desert used in this work.

### Lysine deserts are widespread among bacteria with pupylation pathway

Next, we aimed to ascertain if the lysine deserts may have already emerged in bacteria that employ a pupylation and proteasome-dependent degradation pathway. We analyzed all available bacteria reference proteomes (for 8881 taxons; Table S1) from the UniProt database to check the fraction of proteins (≥150 aa) possessing a lysine desert min. 50%/150 aa in each proteome separately and averaged the results across different taxonomic classes. Interestingly, *Actinobacteria* possess the most proteins with lysine desert min. 50% in their proteomes (Fig 2A), and the same tendency is preserved for lysine desert min. 150 aa (Fig S1A). We also compared proteomes of the most studied bacteria taxons belonging to different taxonomic classes - *M. tuberculosis* H37Rv (virulent), *M. smegmatis*, *C. glutamicum*, *S. ceolicolor*, *L. ferrooxidans*, *B. subtilis*, and *E. coli*. From each aforementioned proteome, we again selected sequences ≥150 aa, and with no more than two predicted transmembrane helices (TMH) using the TMHMM-2.0 software^41, 42^; this condition was applied to exclude proteins with multiple transmembrane regions as they would introduce bias due to their reduced frequencies of polar residues^43^. Applying these criteria resulted in analyzing 58-71% of sequences (see Table 1 in Materials and Methods). Again, the most lysine desert-rich proteomes belonged to the *Actinobacteria* phylum that utilizes the pupylation route for protein breakdown, regardless of the applied lysine desert definition (Fig 2B, Fig S1B). It is noteworthy that the number of sequences with lysine deserts min. 50% was approx. 3- to 4-fold higher in taxons that encode and use the proteasome for regulated degradation of pupylated proteins (25.7% for *M. tuberculosis*, 26.2% for *M. smegmatis* and 35.4% for *S. ceolicolor*) than those that use Pup- modifications but lack proteasomal subunit genes (9.4% for *C. glutamicum*).

**Table 1.** Summary of selected bacteria proteomes for lysine desert analysis.

|  | UniProt proteome ID | Total sequences in proteome | Fraction of filtered<sup>a</sup> sequences |
| --- | --- | --- | --- |
| *M. tuberculosis* H37Rv | UP000001584 | 3995 | 69.54 |
| *M. smegmatis* MC2 155 | UP000000757 | 6602 | 70.72 |
| *C. glutamicum* ATCC 13032 | UP000000582 | 3093 | 64.24 |
| *S. coelicolor* A3(2) | UP000001973 | 8034 | 69.88 |
| *L. ferrooxidans* C2-3 | UP000007382 | 2413 | 65.98 |
| *E. coli* O157:H7 | UP000000558 | 5062 | 62.96 |
| *B. subtilis* subsp. *subtilis* str. 168 | UP000001570 | 4260 | 58.05 |

<sup>a</sup> Excluding sequences <150 aa and with the predicted number of TMH >2.

**Figure 2.**
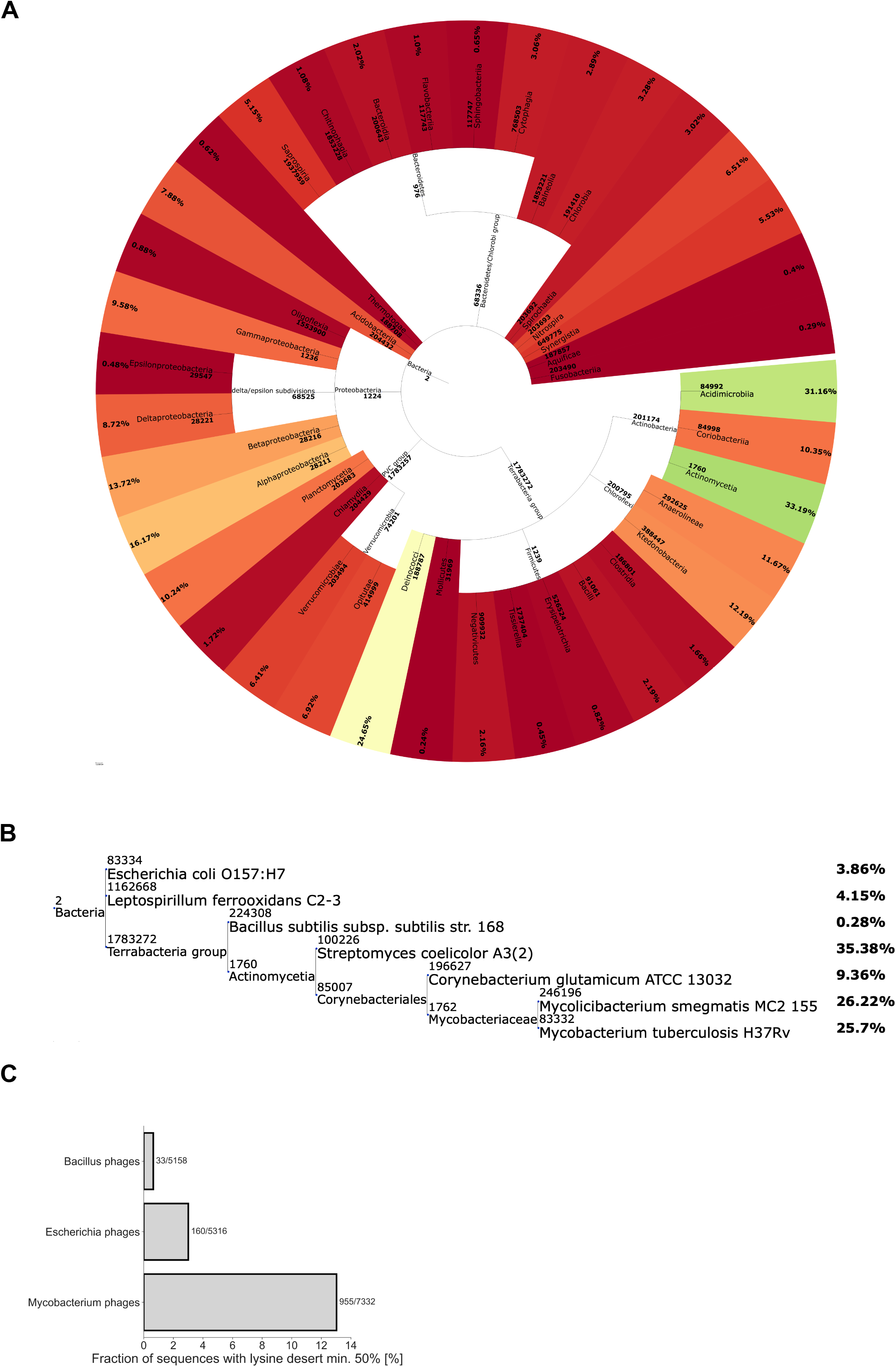
Lysine deserts in bacteria are most prevalent in *Actinobacteria* and their phages. **(A)** Phylogenetic tree of different bacteria classes with a calculated average protein fraction with lysine desert min. 50% in the proteomes of their member taxons. Only classes with at least 10 member taxons were considered. The color gradient corresponds to the min-max normalization where red denotes classes with the lowest average fraction of desert min. 50% and green with the highest. **(B)** Phylogenetic tree of selected bacteria taxons from distinct classes with a calculated fraction of proteins with lysine desert min. 50% in their proteomes. **(C)** Bar plot presenting the fraction of lysine desert min. 50% in pan proteome of selected phages’ groups. The number of sequences with lysine desert and the total number of analyzed sequences are indicated to the right of each bar.

To further investigate the possible linkage of pupylation and lysine desert occurrence, we retrieved information on identified pupylated proteins (pupylomes) of *M. tuberculosis*, *M. smegmatis,* and *C. glutamicum* from the PupDB database^44^, which gathers data from four large-scale proteomics studies. Interestingly, only about 1% of these bacterial species’ whole proteomes (unfiltered in any way) undergo pupylation (Table S2). Similarly as before, we selected sequences ≥150 aa with no more than two predicted TMH among pupylated and non- pupylated proteins of the taxons mentioned above (as non-pupylated we considered all proteins in a proteome except those reported in the PupDB) and searched for lysine deserts among them. We noted that the fraction of sequences with lysine desert min. 50% among pupylated proteins was 1.52% for *M. smegmatis*, 2.56% for *C. glutamicum*, and 8.16% for *M. tuberculosis* (Table S3). The presence of proteins with lysine desert min. 150 aa was also much more prevalent in *M. tuberculosis* (Table S4). Notably, this trend was not related to the average length of filtered sequences in both analyses. In line with the previous results, the fraction of sequences with lysine desert among non-pupylated proteins was approx. 3- (lysine desert min. 50%) to 4- (lysine desert min. 150 aa) fold higher in bacteria employing proteasome to protein turnover (Table S5, Table S6). This observation strongly suggests an as-yet-unidentified mechanism of Pup avoidance on lysines and, thus, proteasomal degradation, similar to yeast examples^35, 36^.

### *Mycobacterium* phages are equipped with lysine desert proteins

Phages are known to be highly specific toward their hosts, co-evolving with them to adopt successful hijacking strategies. We, therefore, decided to inspect whether the difference in lysine desert quantity between *Actinobacteria* and other bacteria phyla also finds reflection in phages specific to different genuses of bacteria. We selected 133 to 527 proteomes of phages specific to *Bacillus*, *Escherichia*, and *Mycobacterium* genuses and clustered them into separate pan proteomes containing 5158 to 7332 sequences ≥150 aa (see Table 2 in Materials and Methods). Interestingly, sequences of *Mycobacterium* phages contain approx. 4- (lysine desert min. 50%) to 5- (lysine desert min. 150 aa) fold more proteins with lysine desert compared to *Escherichia* phages and from approx. 12- (lysine desert min. 150 aa) to 22- (lysine desert min. 50%) fold more proteins with lysine desert compared to *Bacillus* phages (Fig 2C, Fig S1C). This observation suggests an adaptive strategy by phages to minimize the pupylation of viral proteins, potentially avoiding their removal by the host cell.

**Table 2.** Summary of proteomes of Mycobacterium, Escherichia and Bacillus phages.

| Phages group | Proteomes number | Filtered<sup>a</sup> proteomes number | Total sequence number in filtered<sup>a</sup> proteomes | Fraction of sequences from filtered<sup>a</sup> proteomes used for cd-hit | Total number of sequences in pan proteome<sup>b</sup> | Median sequence length in pan proteome<sup>b</sup> |
| --- | --- | --- | --- | --- | --- | --- |
| *Bacillus* | 149 | 133 | 18659 | 40.33 | 5158 | 256 |
| *Escherichia* | 263 | 245 | 22931 | 44.41 | 5316 | 260.5 |
| *Mycobacterium* | 527 | 527 | 53012 | 36.93 | 7332 | 271 |

<sup>a</sup> Excluding outlier proteomes and with <40 sequences.
<sup>b</sup> After running cd-hit.

### Lysine deserts appear with increasing organismal complexity in eukaryotes

We performed a similar screen for the presence of proteins with a lysine desert among the proteomes of five model eukaryotic organisms: S*. cerevisiae*, *C. elegans*, *D. melanogaster*, *M. musculus*, and *H. sapiens*. Again, we excluded very short sequences (<150 aa) with more than two TMH predicted by the TMHMM-2.0 software; hereafter, we refer to those as filtered proteomes. This filtering procedure resulted in the analysis of 66-77% of the sequences (see Table 3 in Materials and Methods). We observed an ascending trend of lysine desert proteome coverage - fractions of lysine desert proteins constituted from 1.04%/2.38% of *S. cerevisiae* proteome to 3.86%/10.5% of *H. sapiens* proteome (lysine desert min. 50%/min. 150 aa, respectively) (Fig 3A, Fig S2A).

**Table 3.**
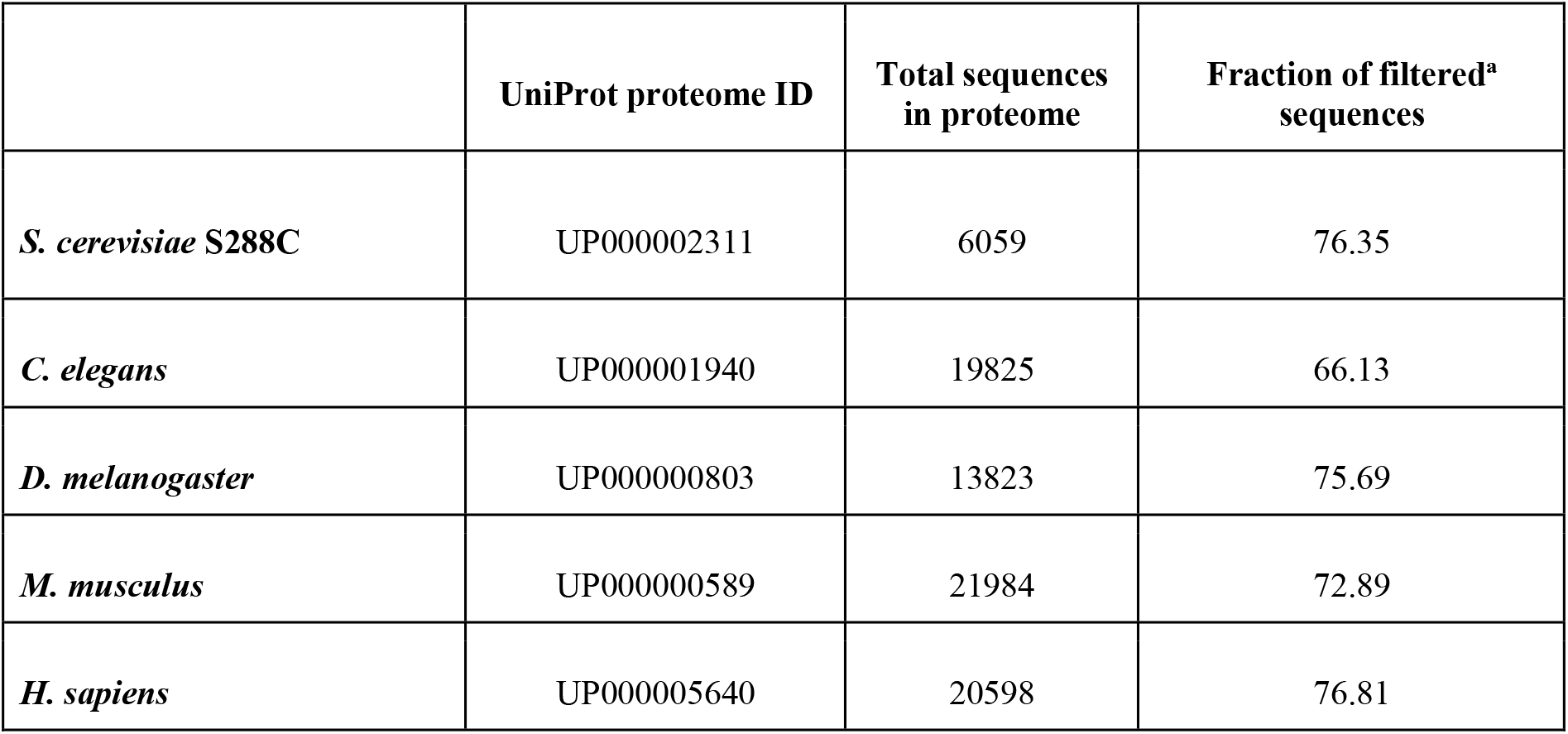
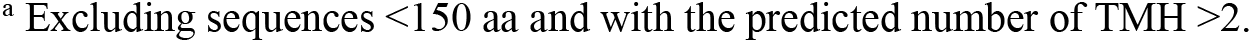
Summary of selected eukaryotic proteomes.

|  | UniProt proteome ID | Total sequences in proteome | Fraction of filtered<sup>a</sup> sequences |
| --- | --- | --- | --- |
| *S. cerevisiae* S288C | UP000002311 | 6059 | 76.35 |
| *C. elegans* | UP000001940 | 19825 | 66.13 |
| *D. melanogaster* | UP000000803 | 13823 | 75.69 |
| *M. musculus* | UP000000589 | 21984 | 72.89 |
| *H. sapiens* | UP000005640 | 20598 | 76.81 |

<sup>a</sup> Excluding sequences <150 aa and with the predicted number of TMH >2.

**Figure 3.**
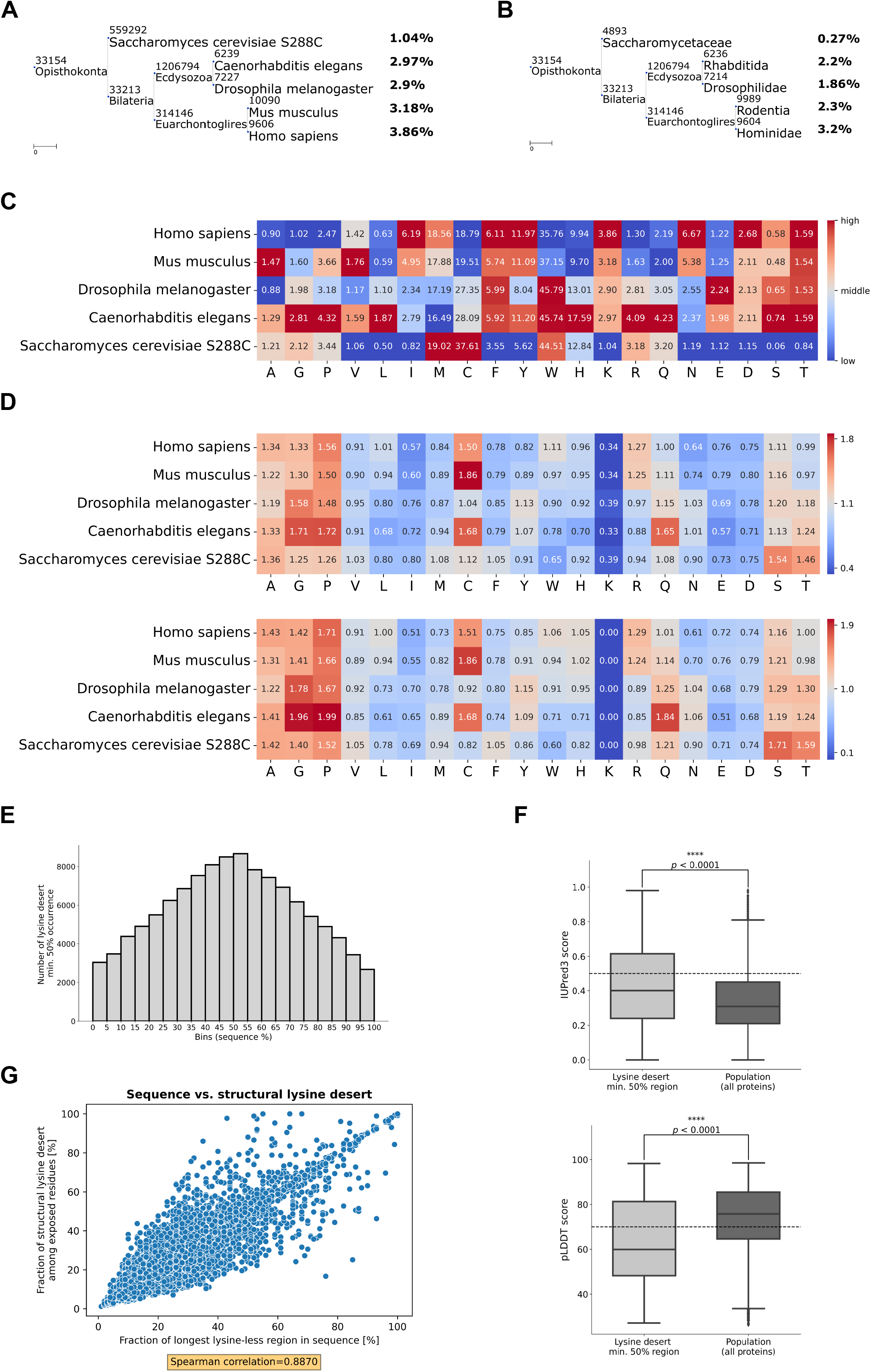
Lysine deserts in eukaryotes ascend with growing organismal complexity. **(A)** Phylogenetic tree of selected eukaryotic model organisms with a calculated protein fraction with lysine desert min. 50% in their proteomes. **(B)** Phylogenetic tree of selected eukaryotic families/order, which include model organisms used in the previous analysis, with calculated fractions of conserved lysine desert min. 50% among their OGs. **(C)** Heatmap of fractions of proteins with desert region min. 50% of each of 20 aa among proteomes of selected eukaryotic model organisms. **(D)** Heatmaps of relative fractions of each amino acid in sequences of selected eukaryotic model organisms with a lysine desert min. 50% normalized to the entire appropriate proteome; value of 1.00 indicates no change. Up - considering whole sequences; down - considering only the lysine desert regions. **(E)** Histogram of distribution of sequence location of lysine desert min. 50% regions in the human proteome. **(F)** Box plots of predicted disorder score of residues constituting lysine desert regions min. 50% only vs. residues of all proteins in the human proteome. Up - sequence-based disorder predictions obtained using the IUPred3 software; higher values indicate a higher disorder probability. Down - structure-based disorder predictions based on the pLDDT values obtained from the AlphaFold2 models of the human proteome; lower values indicate a higher probability of disorder. Disorder cut-offs proposed by the authors of the methods are marked with dashed lines. The stars denote the significance levels per two-tailed *p*-value obtained from the Mann-Whitney U rank test. **(G)** Scatter plot showing co-occurrence of sequence and structural lysine deserts in human proteome based on the AlphaFold2 models. The Spearman rank-order correlation coefficient is denoted below the plot.

Next, we wanted to assess whether the lysine desert regions are conserved among closely related orthologs. We performed a similar analysis of lysine desert coverage within all available Orthologous Groups (OGs; each OG contains sequences of analogous proteins) from the eggNOG5 database^45^ of *Saccharomycetaceae*, *Rhabditida*, *Drosophilidae*, *Rodentia*, and *Hominidae*, which include model organisms used in the above described analysis. As previously, we excluded OGs of short, transmembrane, or unrepresentative proteins (see Materials and Methods and Table 4 for details). Again, we noticed that with the increase of organismal complexity, fractions of OGs with conserved lysine desert proteins were ascending, constituting 0.27%/1.04% of OGs in *Saccharomycetaceae* to 3.2%/9.72% in *Hominidae* (lysine desert min. 50%/min. 150 aa, respectively) (Fig 3B, Fig S2B). These results indicate that lysine desert fractions expand with increasing organismal complexity, and these regions are conserved in homologous proteins, indicating their potential functional involvement.

**Table 4.**
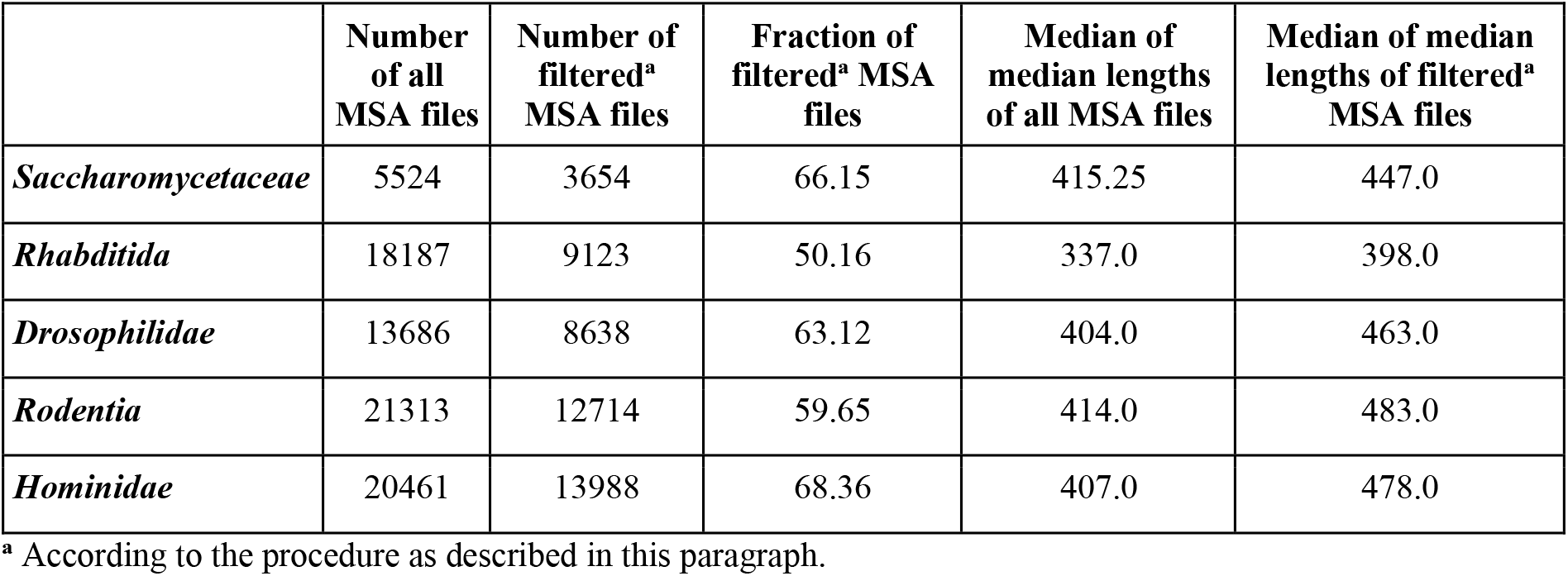
Summary of MSA files of OGs of Saccharomycetaceae, Rhabditida, Drosophilidae, Rodentia, and Hominidae.

We also wanted to assess the uniqueness of the observed tendency of lysine desert ascendance with regard to the amino acid whose absence establishes the desert region. For this purpose, we analogously searched for regions devoid of each of the 20 amino acids constituting min. 50% of the sequence or min. 150 aa in the filtered proteomes of *S. cerevisiae*, *C. elegans*, *D. melanogaster*, *M. musculus*, and *H. sapiens*. We detected a similar gradient trend in isoleucine, asparagine, and aspartic acid for the desert min. 50% (Fig 3C) and additionally in tyrosine and threonine for the desert min. 150 aa (Fig S2C). Interestingly, there was no similar evolutionary ascendance of arginine or histidine desert regions, although these residues, similarly to lysine, yield a positive charge; in fact, we detected a reversed trend for arginine.

Next, we focused on deciphering whether lysine desert proteins are enriched or depleted in particular amino acid(s). We calculated the relative frequencies (each amino acid was normalized by its frequency in the population of a given filtered proteome) of 20 amino acids among the lysine desert proteins and their lysine desert region only in the filtered proteomes of *S. cerevisiae*, *C. elegans*, *D. melanogaster*, *M. musculus*, and *H. sapiens*. We noted that arginine was moderately enriched in *M. musculus* and *H. sapiens* among proteins with lysine desert min. 50% but not min. 150 aa. This may indicate that lysine-less regions could compensate for the lack of lysine with another positively charged amino acid, arginine.We also observed that proteins with lysine desert min. 150 aa and lysine desert min. 50% are enriched (19% - 116%) in alanine, glycine (except for *S. cerevisiae*), and proline in all analyzed eukaryotic model organisms (Fig 3D, Fig S2D), which may point to the occurrence of low complexity regions within them^46^.

As lysine desert proteins show features characteristic of low-complexity regions we aimed to investigate their structural features in the human proteome. First, to obtain a picture of the preferred location of the lysine desert regions, we analyzed their distribution among all protein sequences (again, when referring to the human proteome, we mean its filtered version as described previously; data on lysine desert occurrence of a given type in each human protein are available in Table S7). Interestingly, regions of lysine desert min. 50% tend to occupy internal parts of the protein, avoiding the N-/C-terminus (Fig 3E). This tendency, albeit not as pronounced, is also evident for lysine desert defined as min. 150 aa (Fig S2E).

Since the lysine deserts of yeast San1 and Slx5 E3s are mainly disordered, we sought to determine the structural status of lysine desert regions throughout the human proteome. For each protein, we predicted its disorder score based on its sequence using the IUPred3 software^47^ and obtained the pLDDT (predicted Local Distance Difference Test) values, which estimate the modeling accuracy of each residue^48^, from the corresponding AlphaFold2 model^49, 50^ (see Materials and Methods). Both techniques have been demonstrated to be the gold standard for forecasting disordered regions^47, 51^. Using a sequence- and a structure-based method, our approach allowed us to get unbiased and consistent results - lysine desert regions, either min. 150 aa or min. 50% are more disordered than the entire human proteome (Fig 3F, Fig S2F).

Since protein structure constitutes the interface of intermolecular interactions and therefore plays an essential role in protein functioning, we aimed to evaluate whether lysine desert regions detected in sequences are also maintained in structures. To this end, we screened AlphaFold2 models of the human proteome^50^ to search for the structural lysine deserts. Of note, we did not use any cut-off values regarding its length as in the case of sequence lysine desert (Fig 1C); rather, we defined it as the most extended, uninterrupted lysine-less region among solvent-exposed residues remaining in contact (see Materials and Methods for the details on our algorithm). Our analysis showed a strong correlation between the length of lysine-less regions in sequence and the structure of the human proteome (its filtered version; we also excluded from the analysis proteins with more than 5% of residues without calculated solvent accessibility values) (Fig 3G). Interestingly, the coverage of residues building structural lysine deserts among residues constituting sequence lysine deserts (in other words, the common residues between sequence/structural lysine desert) varies, with most pronounced cases where the longest lysine-less region constituting over 40% of the sequence does not overlap with the structural lysine desert in that protein (Fig S2G). This indicates how important it is not to overlook the structures, as the information encoded in the sole sequence of a protein may provide a misinterpretation of data. Information on the length and residues building the three most extended structural lysine deserts, tabulated with information about the longest lysine-less regions present in the sequence, can be found in Table S8.

### Most evolutionarily conserved lysine desert proteins operate within the UPS pathway and tend to group their lysines into clusters

Yeast E3 ligases remain the only identified functional lysine deserts^35, 36^. Therefore, we decided to establish if lysine-free sequences are exclusive to the UPS pathway. To this end, we analyzed the function of lysine desert proteins and their evolutionary conservation among organisms with comparable and vastly different levels of complexity. First, we checked which proteins of *S. cerevisiae*, the simplest eukaryotes studied here, possess conserved lysine desert min. 50% among their orthologs in the *Saccharomycetaceae* family. We used the same set of OGs from *Saccharomycetaceae* and the methodology of defining a conserved lysine desert as previously described (see Materials and Methods). As the criteria for recognizing OGs as conserved lysine desert-containing were very stringent, we found 10 such cases for the following proteins: Cue1, Mix17, Rad23, San1, Slx5, Tif6, Tir3, and YMR295C (gene names provided for *S. cerevisiae*) (see Table S9; summaries of OGs with conserved lysine desert min. 150 or 50% in *Saccharomycetaceae*, *Rhabditida*, *Drosophilidae*, *Rodentia*, and *Hominidae* are available at https://github.com/n-szulc/lysine_deserts52). Notably, six of the listed - Cue1, Dsk2, Hlj1, Rad23, San1, and Slx5, regulate protein turnover (Fig 4A).

**Figure 4.**
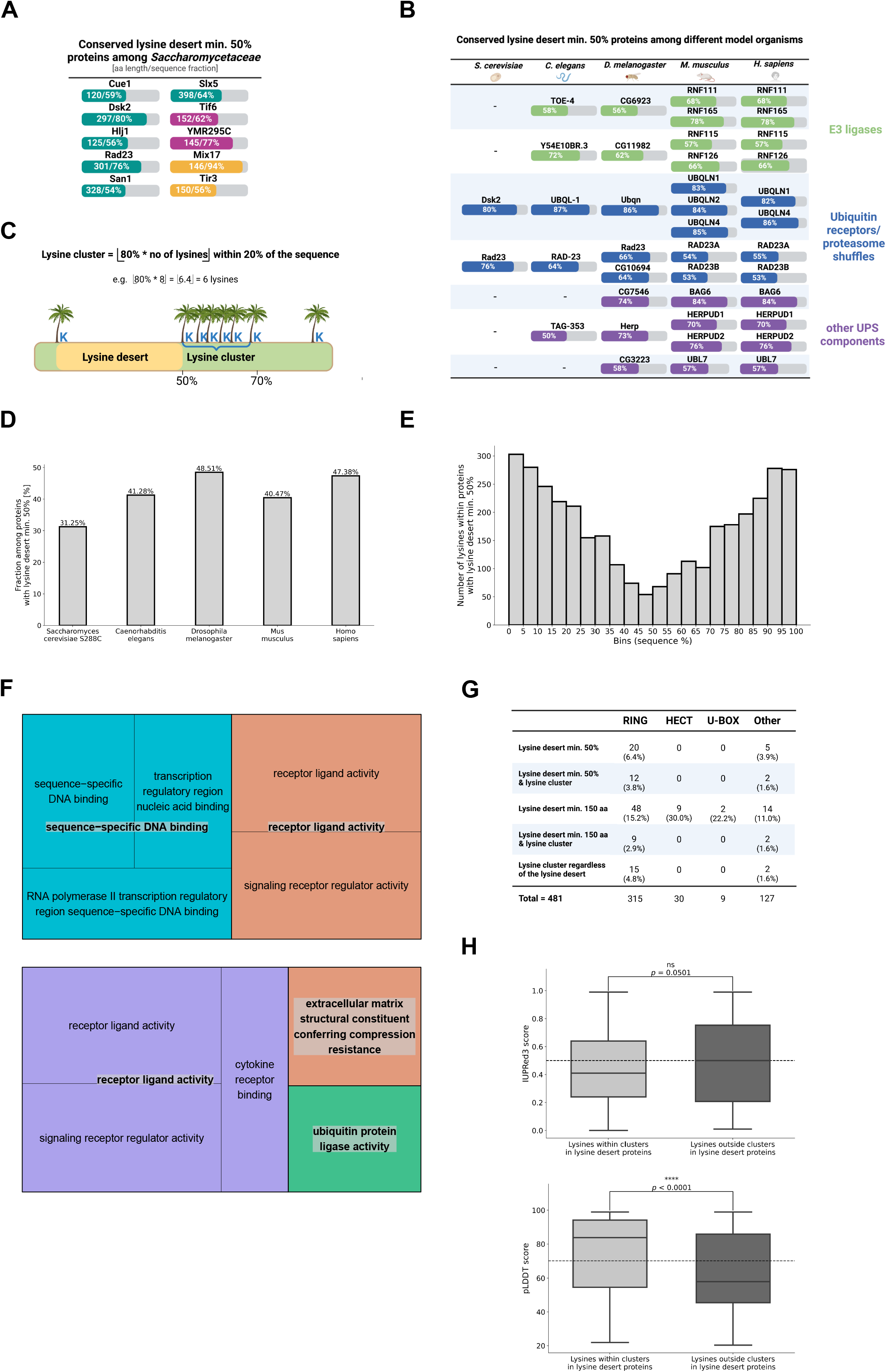
Lysine deserts are conserved among UPS proteins and often co-occur with a lysine cluster. **(A)** Conserved lysine desert min. 50%-containing proteins among the *Saccharomycetaceae* family. Gene names are provided for *S. cerevisiae.* Nominal values of the lysine desert’s length and sequence fraction in each protein are denoted on the bars. Teal - proteins associated with protein turnover; magenta - ribosomal proteins; yellow - other proteins. **(B)** Proteins with the most evolutionarily conserved lysine deserts. The selection criteria were as follows: lysine desert min. 50% conserved in OGs from *Drosophilidae*, *Rodentia,* and *Hominidae* (in *Saccharomycetaceae* and *Rhabditida*, OGs were either absent or, if present, conserved lysine desert min. 50% also needed to occur). Each row represents one vector set of analogous OGs. For clarity, fractions of lysine desert regions in proteins from the model organism representative of the aforementioned families/order are shown. **(C)** Definition of a lysine cluster. **(D)** Bar plot of the fraction of lysine clusters’ occurrence among proteins with lysine desert min. 50% from selected model organisms. Exact values are denoted above each bar. **(E)** GO molecular function terms found to be associated with proteins with lysine desert min. 50% (up) and with proteins with lysine desert min. 50% co-occurring with a lysine cluster (down). **(F)** Histogram of distribution of sequence location of lysines within proteins with lysine desert min. 50% from the human proteome. **(G)** Summary of co-occurrence of lysine deserts min. 150 aa and min. 50% and lysine clusters among human E3s. **(H)** Box plots of predicted disorder score of lysine residues belonging to a lysine cluster among proteins with lysine desert min. 150 aa and/or min. 50% vs. lysine residues, which do not form a lysine cluster among proteins with lysine desert min. 150 aa and/or min. 50% in the human proteome. Up - sequence-based disorder predictions obtained using the IUPred3 software; higher values indicate a higher disorder probability. Down - structure-based disorder predictions based on the pLDDT values obtained from the AlphaFold2 models of the human proteome; lower values indicate a higher probability of disorder. Disorder cut-offs proposed by the method’s authors are marked with dashed lines. The stars denote the significance levels per two-tailed *p*-value obtained from the Mann-Whitney U rank test.

Next, we aimed to compare the evolutionary conservation of lysine deserts min. 50% between analogous OGs of different eukaryotic families/orders (e.g., whether protein X with a conserved lysine desert in a given OG of *Saccharomycetaceae* also has a conserved lysine desert in an analogous OG of *Hominidae;* we term such analogous OGs as vectors). For this analysis, we examined the occurrence of conserved lysine deserts in the vectors of respective OGs (see Materials and Methods for details; complete results are available at https://github.com/n-szulc/lysine_deserts). We applied stringent criteria for the presence of lysine desert min. 50% conserved in OGs from *Drosophilidae*, *Rodentia,* and *Hominidae* among 4282 vectors of known orthologs from these families/order (for *Saccharomycetaceae* and *Rhabditida*, OGs either could be absent or, if present, conserved lysine desert min. 50% also needed to occur). We again found proteins responsible for protein degradation (7 out of 9 vectors; see Table S10) - E3s, ubiquitin-adaptor proteins, and other proteostasis components (Fig 4B).

Interestingly, during our analyses we noticed that lysines in some lysine desert proteins (e.g., RNF126, RNF165, ubiquilins, HERPUD1, HERPUD2) occur within a relatively limited part of the sequence. We reasoned that such clustering of lysines while maintaining a large lysine desert region might have a regulatory role in turnover control. We estimated the frequency of lysine clusters within the lysine desert proteins to examine how this tendency relates to our global studies of selected model organisms’ proteomes (again, their filtered versions as previously described). We arbitrarily defined a lysine cluster as ⌊80% * total number of lysines in sequence⌋ within 20% of sequence; applies only to proteins with min. two lysine residues (Fig 4C). Over 47% of human lysine desert min. 50% proteins possess a lysine cluster, and other model organisms also show high (31-49%) such co-occurrence (Fig 4D).

However, this trend is much less pronounced for lysine desert min. 150 aa, where lysine cluster occurs in 10-16% of such proteins from the analyzed model organisms (Fig S3A). Relating to the global filtered proteomes, lysine clusters occur in 0.8-3.3% of proteins (Fig S3B), which indicates that the size of the desert may correspond with the existence of the lysine cluster. We also analyzed lysines’ distribution among sequences of human proteins possessing lysine desert regions. Although lysine deserts favor internal parts of the protein (Fig 3E, Fig S2E), lysines show a much more bimodal distribution with peaks at N-/C-terminus (Fig 4E, Fig S3C). Even among proteins with lysine desert min. 150 aa, which have a much flatter distribution of their lysine-less regions (Fig S2E), there is a visible tendency of lysines to localize at the C-terminus (Fig S3C).

To evaluate molecular functions associated with lysine deserts and lysine clusters, we performed Gene Ontology (GO)-based overrepresentation analysis of genes derived from sets of human proteins with lysine desert min. 50%. We observed that molecular functions over- represented among proteins possessing lysine desert min. 50% are broadly defined as DNA-binding and receptor-ligand signaling (Fig 4F, upper panel), indicating the diverse roles of such proteins. Similarly, for lysine desert defined as min. 150 aa, the molecular functions are broadly annotated (Fig S3D). Noteworthy, for proteins where a lysine desert min. 50% co-occurs with a lysine cluster, ubiquitin ligase activity is over-represented (Fig 4F, lower panel), which may point to the functionality of this lysine division in the context of a vast lysine desert, especially within the ubiquitin-signaling components. The abovementioned results prompted us to look closely at the occurrence of lysine deserts and lysine clusters among E3 ligases. Since there are inconsistencies in the literature regarding the number of E3 ligases in the human proteome, we manually curated a list of 481 human proteins associated with ubiquitin ligase activity and their type annotation (Table S11). We detected that 25 of them possess a lysine desert min. 50%, and within them, 14 also have a lysine cluster (Fig 4G). Interestingly, when we analyzed E3s by their types, it was remarkable that the lysine desert and lysine cluster are typical features of the RING E3 ligases - 12 out of 14 E3s with both lysine desert min. 50% and a lysine cluster were of this type; a similar trend also occurs for lysine desert min. 150 aa (Fig 4G). In addition, lysine-deficient regions are primarily found in the disordered regions of the human RING E3s (Fig S3E, F).

Similarly, as for lysine deserts, we wanted to check whether lysine clusters co-occurring with lysine deserts tend to locate in disordered regions of proteins. Using the same approach as previously, we noted that lysines of lysine clusters are more structured than other lysines among human proteins with lysine desert (Fig 4H). Intriguingly, many human E3s, such as aforementioned highly conserved RNF126 and RNF165, but also other conserved among mammals, e.g., RNF6 (lysine desert of 527 aa, constituting 77% of the sequence), RNF12/RLIM (lysine desert of 477 aa, constituting 76% of the sequence), or RNF44 (lysine desert of 350 aa, constituting 81% of the sequence) aggregate their lysine residues within the RING domain, which interacts with E2s. Presumably, owing to the lysine clusters in the functional domains, E3 can undergo precise auto-ubiquitination without modification in the vast lysine desert region, which i.e., could affect substrate binding.

### CRL substrate receptors are deprived of lysines in the course of evolution

The restricted availability of orthologous sequences from evolutionarily distant organisms (such as invertebrates and mammals) poses a challenge for comparative analyses. Moreover, searching only for long lysine-less regions may prevent the detection of proteins that lost lysines during evolution but do not have lysine-free stretches long enough to surpass the predetermined threshold (e.g., the threshold for considering protein as containing a lysine desert min. 50%). For these reasons, we searched for E3 ligases in the filtered human proteome possessing max five lysines (Table S7), excluding those described in the UniProt database as membrane-bound (even single-pass). We tabulated the obtained lysine-poor E3s with their distant vertebrate orthologs from *D. rerio*, *X. tropicalis*, and *G. gallus,* as well as from more closely related *M. musculus,* based on the information from the eggNOG5 and Xenbase^53^ databases (Table S12). Out of 14 such human E3 ligases, eight showed significant lysine “desertification” (from 2- up to a 15-fold decrease of lysines) in the course of evolution (Fig 5A). Notably, the majority (seven out of eight) of such E3s act as substrate receptors subunits of CRLs. While operating within the complexes, these proteins risk being labeled by ubiquitin during binding client proteins and bringing them into the vicinity of the E2 enzyme for ubiquitination.

**Figure 5.**
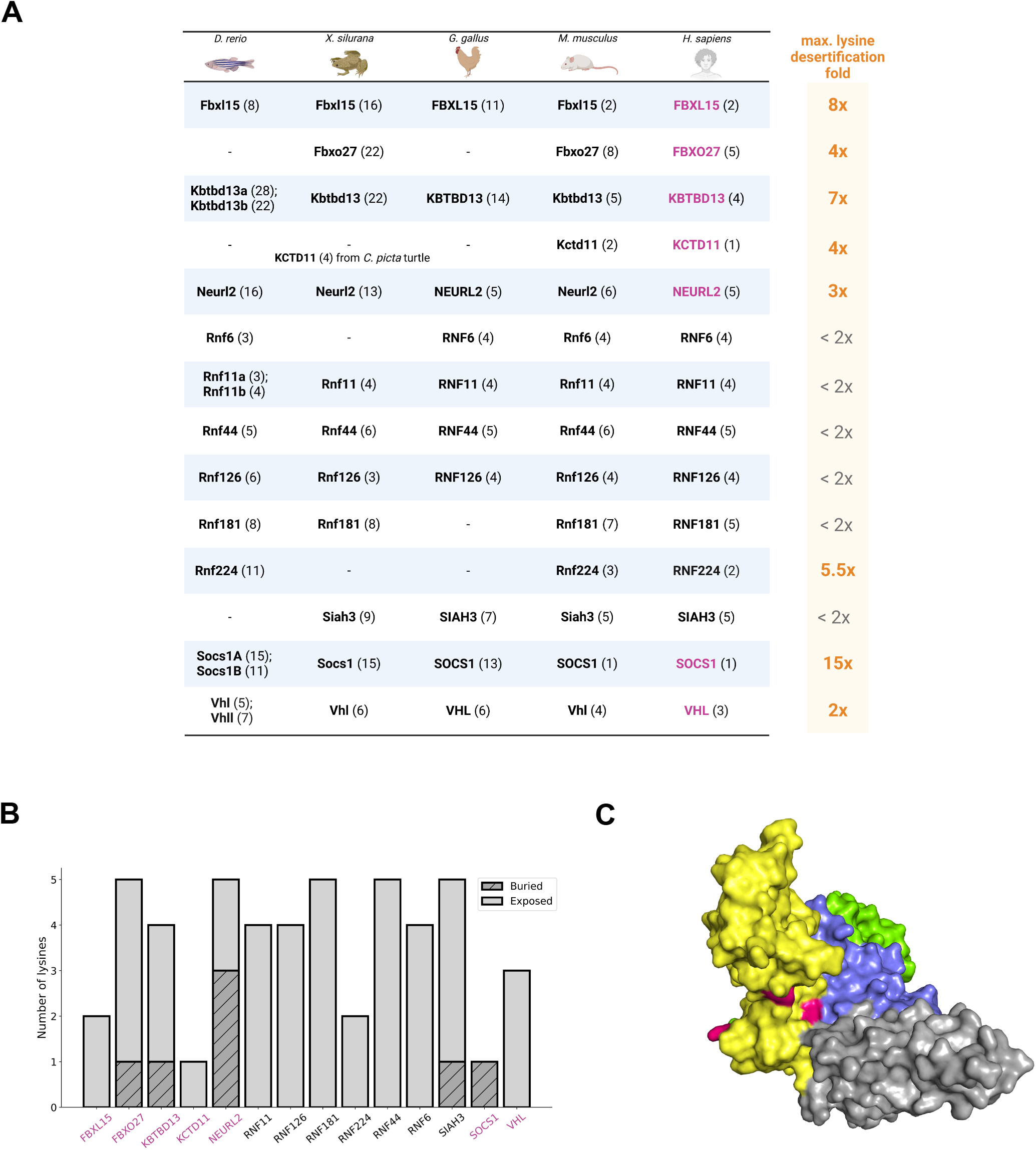
Human CRL substrate receptors are evolutionarily deprived of lysines and restrict solvent accessibility for the remaining ones. (A) Human E3 ligases possessing max. five lysines and their orthologs from *D. rerio*, *X. silurana, G. gallus,* and *M. musculus*. A dash marks unknown orthologs; due to genome duplication in *D. rerio* its orthologs are separated with a semicolon. Due to unknown orthologs from *D. rerio*, *X. silurana,* and *G. gallus* of KCTD11, the number of lysines in its ortholog from *C. picta* turtle was reported. The total number of lysines for each protein is denoted in brackets. Human CRL substrate receptors are marked in magenta; maximum evolutionary lysine desertification is indicated to the right. (B) Bar plot of the number of exposed and buried lysines in human E3 ligases possessing max. five lysines. Relative solvent accessibility, based on which residue is classified as exposed or buried, was calculated using the DSSP program and the Sander method (see Materials and Methods for details). Residues with RSA >0.2 are considered as solvent-exposed. Human CRL substrate receptors are marked in magenta. (C) VHL in complex with cullin 2 (PDB ID: 4WQO). Color codes explanation: yellow - VHL, green - elongin B, blue - elongin C, grey - cullin 2, magenta - lysine residues (three in total) of VHL. Visualized in the PyMOL software (Schrödinger) (v. 2.5.0).

Since lysine is one of the most solvent-accessible amino acids^54^, we next aimed to determine if the few lysines of the abovementioned CRL substrate receptors subunits are also exposed to solvent. We performed the solvent accessibility analysis precisely the same as for the structural lysine desert search (see Materials and Methods for details) based on the AlphaFold2 models of selected lysine-poor CRLs substrate receptors subunits, as their experimental structures, except for VHL, remain unsolved. Interestingly, not all their lysines are solvent accessible, making them even less prone to lysine-dependent ubiquitination, with the most extreme case of SOCS1, which decreased its lysines’ content 15-fold and possesses only one but buried lysine (Fig 5B, Table S13). VHL complex-free monomer has all its lysines exposed (Fig 5B), but when it associates with the elongin B, elongin C, and cullin 2, access to one or two lysines may be restricted (Fig 5C), likely making them inaccessible for any modifications. Moreover, the burial of lysines within the complex might be exploited by other CRL substrate receptors; therefore, their lysine solvent accessibility could differ from the one calculated for their models of complex-free monomers. The above analyses may indicate an evolutionary pressure to limit the ubiquitination of these CRL substrate receptors.

### Lysine desert human proteins, SOCS1 and VHL, undergo non-canonical ubiquitination and ubiquitin-independent proteasomal degradation

To assess the involvement of the UPS in the regulation of CRL substrate receptors subunits, we measured the ubiquitination of SOCS1 and VHL, as well as their all-lysine-deficient variants (VHL K159R, K171R, K196R; SOCS1 K118R), in living HEK293 cells. To this end, we utilized the NanoBRET technology (Promega), which involved the transient expression of NanoLuc fusions (at the C- or N- terminus) of SOCS1 and VHL variants and an N-terminal HaloTag-ubiquitin fusion, followed by the NanoBRET assay, the output of which is a signal that increases proportionally to the degree of ubiquitination. Both wild-type SOCS1 and VHL, as well as their lysine-free variants, exhibited comparable ubiquitination levels. Interestingly, inhibition of the proteasome achieved by treating the cells with MG132 inhibitor (cell viability assessed with CellTiter-Glo following treatment was unaltered; see Fig S4A, B) did not result in the accumulation of ubiquitinated SOCS1 and VHL variants, which may suggest that they undergo non-lysine ubiquitination, which does not contribute to proteasome-dependent degradation (Fig 6A, B).

**Figure 6.**
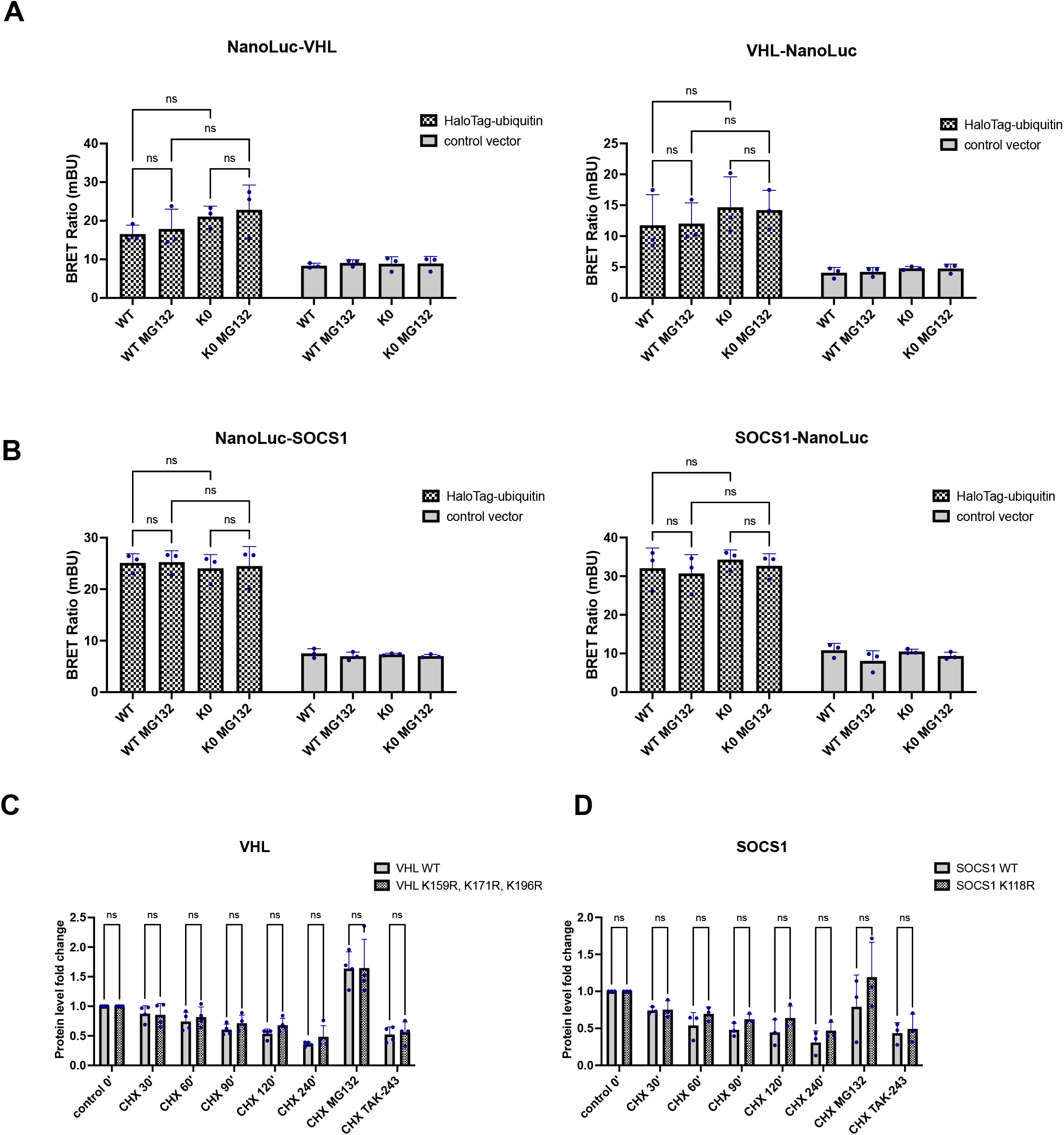
Lysine deserts provide a platform for non-lysine ubiquitination and can be degraded by proteasomes independent of ubiquitination. (A-B) VHL and SOCS1 wild-type and their lysine-less variants’ ubiquitination measured by NanoBRET assay with transient expression of the HaloTag-ubiquitin or the control vector pHTN HaloTag CMV-neo and NanoLuc-tagged (at its N-/C-terminus as indicated) protein of interest. Where specified, cells were also co-treated with 20 µM MG132 proteasome inhibitor for 2 hours. K0 indicates a lysine-less variant (K159R, K171R, K196R for VHL and K118R for SOCS1). Ubiquitination levels were measured as described in Materials and Methods. Error bars represent the standard deviation of the mean of three independent biological replicates, dots represent the biological replicates. Each biological replicate is a mean of three technical replicates. The significance levels obtained from Tukey’s multiple comparisons test are indicated for the compared conditions (ns - not significant). Analyzed and visualized in the GraphPad Prism 9. (C-D) VHL and SOCS1 wild-type and their lysine-less variants’ turnover measured by CHX assay with transient expression of the HiBiT-tagged protein of interest and treatment with 50 µg/ml CHX for the indicated time. Where specified, cells were co-treated with 20 µM MG132 proteasome inhibitor for 4 hours or pre-treated with 1 µM TAK-243 E1 inhibitor for 1 hour before CHX was added for the next 4 hours. Protein levels were measured and normalized to the number of living cells as described in Materials and Methods; chase assays were normalized to the corresponding measurement for each protein variant from control time 0’. Error bars represent the standard deviation of the mean of independent biological replicates (four for VHL and three for SOCS1 experiments), dots represent the biological replicates. Each biological replicate is a mean of three technical replicates. The significance levels obtained from Šídák’s multiple comparisons test are indicated for the compared conditions (ns - not significant). Analyzed and visualized in the GraphPad Prism 9.

To determine the turnover of SOCS1 and VHL and their lysine-free variants, we carried out cycloheximide chase assays (CHX). Here, we assessed the levels of SOCS1 and VHL variants tagged with High BiT (HiBiT, 11 amino acid tag; Promega) in HEK293 cells and detected their levels with a reagent containing the complementary peptide Large BiT (LgBiT; 17.6 kDa; Promega), which binds to HiBiT with high affinity generating luminescence. This provides quantitative protein abundance measurements in a linear dynamic range. Noteworthy, there is no risk of introducing additional ubiquitination sites when tagging proteins with HiBiT, as the HiBiT tag was validated as not prone to ubiquitination (information from the Promega R&D Department). We did not detect significant differences in the stability of SOCS1 and VHL variants during the CHX chase. With this assay, we also determined if inhibition of ubiquitination via the E1 inhibitor (TAK-243)^55^, or the proteasome inhibitor (MG132) impacts the wild-type and lysine-free SOCS1 and VHL degradation rates. Interestingly, E1 inhibition did not affect the stability of these CRL receptors, in contrast to the proteasome inhibition, where there was a significant accumulation of SOCS1 and VHL wt and lysine-free variants (Fig 6C, D). CHX assays and NanoBRET ubiquitination measurements revealed that the lysine deserts in SOCS1 and VHL do not prevent their ubiquitination but enable non-canonical ubiquitin labeling. In addition, these modifications are rather regulatory and do not increase proteasome targeting. Nevertheless, irrespective of ubiquitination, SOCS1, and VHL are degraded by the proteasome.

### Lysine deserts do not correlate with human proteins half-live

Since our experiments demonstrated that lysine deserts do not reduce the possibility of proteasomal degradation, we investigated whether there is a correlation between the half-life and the presence of lysine deserts in cell-specific human proteomes. To this end, we obtained protein turnover datasets from two large-scale proteomic studies^56, 57^. The dataset prepared by Mathieson and colleagues provided information on the turnover of 3555 - 4653 human proteins in primary cells, namely B cells, NK cells, hepatocytes, and monocytes, while Li and colleagues measured half-lives of 1428 - 1904 short-lived human proteins in four cell lines: U2OS, HEK293T, HCT116, and RPE1 (see Materials and Methods for a summary of the number of analyzed proteins in each cell type). We found no correlation between the length of the lysine desert and the length of protein half-live in any dataset, regardless of whether we compare the nominal length of the lysine desert region or its fraction of the protein sequence (Fig 7, Fig S5). Together with ubiquitination and turnover analyses of SOCS1 and VHL, these global analyses suggest that the presence of lysine deserts does not contribute to the half-life of equipped human proteins, as they may be subject to efficient non-canonical regulation by the UPS system.

**Figure 7.**
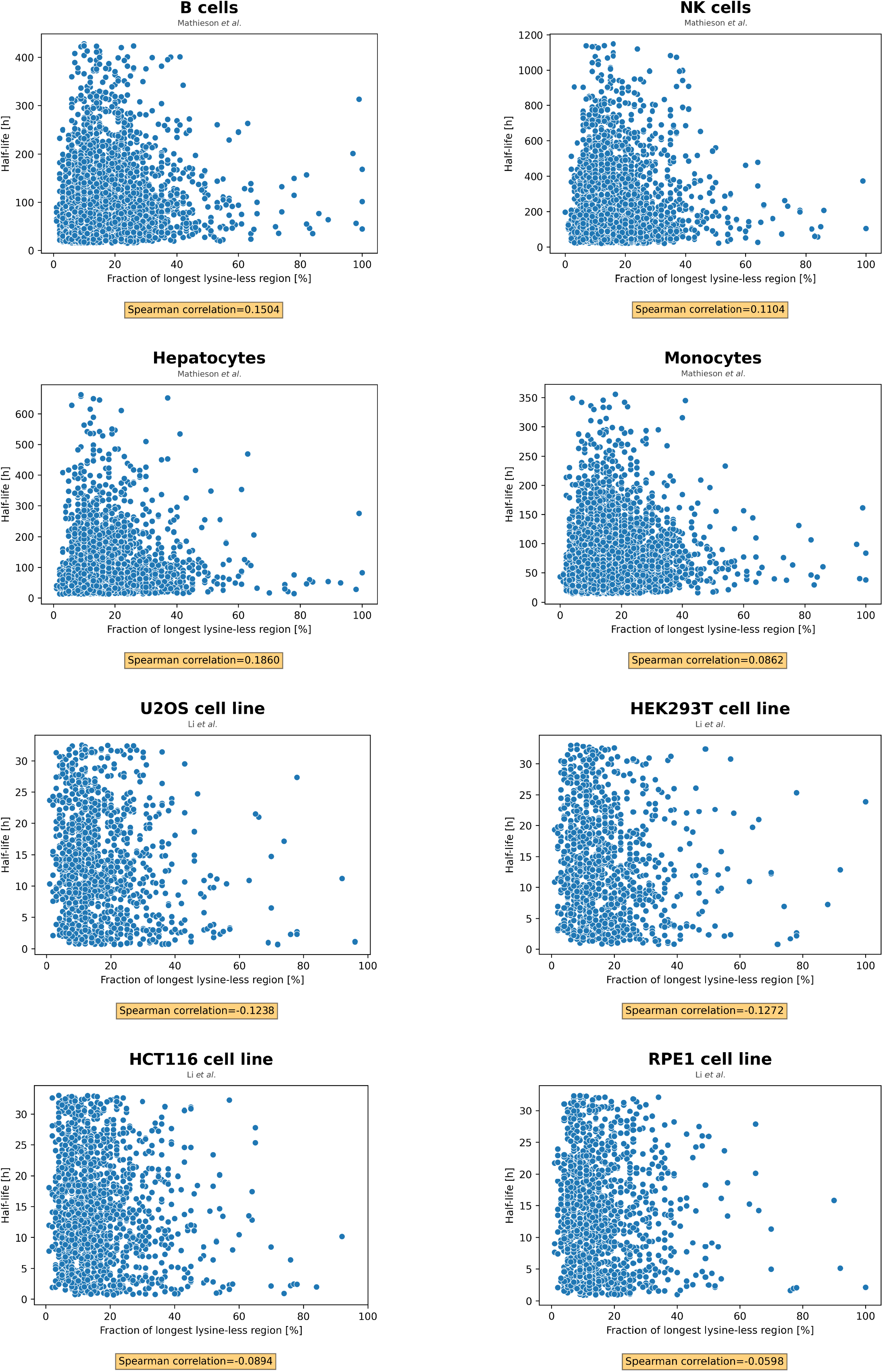
Lysine deserts do not correlate with protein half-life. Scatter plots showing a lack of correlation between the length of the lysine-less region and protein half-life in eight different cell types from two large-scale proteomic studies of human proteins. The Spearman rank-order correlation coefficient is denoted below each plot.

## Discussion

A functional components of the UPS exposed to ubiquitination should be equipped with mechanisms to prevent accidental proteolytic destruction. Selective pressure to avoid lysine residues, especially in intrinsically disordered regions, may underpin a strategy to avoid redundant ubiquitination of ubiquitin-proteasome components. To determine this possibility, we designed and implemented a bioinformatic pipeline to quantitatively search for lysine-free regions and investigate their evolution and functional roles among orthologous prokaryotic and eukaryotic taxa.

We noted that the abundance of lysine deserts in *Actinobacteria* is most prevalent among species that possess proteasomes and utilize a pupylation pathway. The Pup-proteasome system (PPS) plays a key role in mycobacterial stress responses^58^. For example, nitrogen starvation or reactive nitrogen species secreted by host macrophages in response to infection by *M. tuberculosis* induces PPS^58, 59^. Perhaps the abundance of lysine deserts in the *M. tuberculosis* proteome promotes the feasible degradation of only specific nitrogen metabolic network components, such as the HrcA repressor of chaperonin, which promote the nitrite reductase NirBD to assimilate nitrogen from nitrate^60^. Furthermore, we found that *Mycobacterium* phages’ sequences contain several to dozens of times more lysine-depleted proteins than phages of bacteria not equipped with the PPS. Possibly this contributes to limiting pupylation and degradation of phages’ proteins enabling more efficient infection or killing of the mycobacterial host exposed to stress conditions.

We performed a similar screen for the presence of proteins with a lysine desert among the proteomes of model eukaryotic organisms and identified many E3s, ubiquitin-adaptor proteins, and other components of the cellular proteostasis system containing long sequence stretches completely devoid of lysine. One example of the latter is the ubiquitin-like (UBL) domain of BAG6. Kampmeyer and colleagues showed that introducing lysine residues into the BAG6 lysine-free sequence leads to increased ubiquitination and proteasomal degradation driven by its associated partner, E3 RNF126^61^, which itself is also an example of a protein with highly conserved lysine desert^62^. We observed that in human E3s, predominantly among the class of the RING ligases, lysine-depleted regions are present primarily in disordered regions (Fig S3D, E). Perhaps avoidance of lysine modification by ubiquitin in these parts is required for their localization, conformation, activity and substrate binding^63, 64^. In addition to the UPS-related proteins, we identified several components of the multisubunit molecular complexes, such as NF-Y and the RNA exosome complex, equipped with an extensive lysine-deficient region. Notably, in NF-Y, only NFYA protein is devoid of lysines, whereas, in the RNA exosome complex, only two of its nine subunits, EXOSC4 and EXOSC6, contain extensive lysine deserts. We speculate that these lysine-deficient proteins may serve as a homing element for the UPS to target other components of the complexes or act as a stable seed-initiating protein (de)complexation.

Interestingly, many lysine desert E3s aggregate their lysine remnants within the RING domain, which interacts with E2. This may indicate pressure for specific auto-ubiquitination in the ordered cluster zone but not in the disordered lysine-free region. The example of RNF12/RLIM, a 624 aa long E3 ligase with a large disordered lysine desert undergoing intensive auto-ubiquitination on the RING-localised lysine cluster while sparing the remaining part responsible for sorting and substrate binding, seems to support this hypothesis^65–67^. Another example of highly conserved lysine desert proteins with lysine clusters are ubiquilins (UBQLN1, UBQLN2, UBQLN4/Ubqn/UBQL-1), which recognize ubiquitinated proteins and guide them to the proteasome^68^. Ubiquilins locate all of their lysines to the N-terminal UBL domain, which involves interaction with proteasome receptors, leaving the UBA domain (ubiquitin-associated domain), responsible for recognizing and binding ubiquitin on client proteins^68, 69^, completely lysine devoid. Therefore, ubiquilins likely do not expose lysines to the cytoplasmic environment when binding to the proteasome, presumably being protected against ubiquitination during functioning. Upon the substrate/partner dissociation, the protein may again become susceptible to ubiquitination. Thus, lysine clusters can be functional regulatory elements of lysine desert E3s and other proteostasis factors.

When searching for the most evolutionarily conserved lysine desert proteins, we noticed a gradient trend in the appearance of lysine deserts in higher eukaryotes and hypothesized that some proteins expand their lysine-depleted region as organisms evolve. The most prominent examples are substrate receptors of the CRLs like VHL. *C. elegans* VHL-1 and *D. melanogaster* Vhl proteins’ lysine-less region fraction is 26% and 31%, respectively, while in contrast, their orthologs in *M. musculus* and *H. sapiens* have lysine deserts stretching for 66% and 74% of their sequences, respectively, suggesting evolutionary pressure on lysine-free region elongation. Besides, the lysine desert region appears in VHL not as a result of sequence lengthening but due to the conversion of lysine present in simpler organisms (worm and zebrafish) to other amino acids (i.e., arginine) in mouse and human homologs. Notably, the lysine-less region in VHL covers the VHL substrate-binding domain^70^. SOCS1, also a CRL substrate receptor, is another example. Human and mouse SOCS1 have only one lysine, while its orthologs in lower organisms possess multiple. The number of lysines in SOCS1 decreased by 14 from zebrafish to mouse/human (no ortholog is known in invertebrates), and most of the lysines were replaced by arginines, which also possess a positive charge but cannot be tagged with ubiquitin. Avoiding lysines in such receptors may reduce their risk of being labelled by ubiquitin when binding client proteins and bringing them close to E2 for modification. Thus, we initially speculated that the expansion of the lysine desert region in mammals is correlated with the development of UPS and redundancy of E2 and E3, providing protection against non-specific degradation on their part and promoting the lifetime of lysine-depleted proteins. However, we did not observe a correlation between the extent of lysine deserts and protein half-life in human protein turnover data sets from proteomics studies (Fig 7, Fig S5). This may indicate that a lysine desert does not generally increase protein stability but only reduces lysine ubiquitination. Correspondingly, lysine desert in proteasome substrate shuttle RAD23A protects against its ubiquitination but does not affect proteasome degradation^62^. This raises the question of whether the UPS can use non-lysine ubiquitination or proteasomal degradation independent of ubiquitination to regulate the lysine-deficient proteome.

To investigate the turnover of lysine desert proteins in relation to non-canonical regulation by the UPS, we conducted studies with human VHL and SOCS1 proteins and their lysine-free variants. Using the NanoBRET technology, we performed measurements of intracellular ubiquitination of VHL and SOCS1 variants in HEK293 cells under normal growth conditions and noted that they are subjected to modification by ubiquitin at non-lysine positions. In a quantitative kinetic degradation assay based on the HiBiT tagging system, we showed that VHL and SOCS1, and their lysine-free variants, displayed similar turnover rates, which did not depend on ubiquitination, but on proteasomal activity. However, this result is not entirely consistent with previous studies. Pozzebon and colleagues showed that the expression of Gam1, an adenoviral protein, induces VHL ubiquitination and degradation that depends on cullin 2 and cullin 5^71^. However, these studies did not characterize the type of ubiquitination nor track VHL turnover in the case of global ubiquitination inhibition by the E1 inhibition. Wu et all suggested that SOCS1 is subject to ubiquitination which is enhanced by proteasome inhibition and regulated by CUEDC2^72^. Yet, their experimental approach, which makes the analysis of non-lysine modification virtually impossible, indicates extensive polyubiquitination of SOCS1, despite only one lysine in this protein. Moreover, SOCS1 pull-down approach used in this study did not exclude the scenario that these modifications involve substrates bound by SOCS1. Thus, the results of both reports do not preclude our conclusions, which are based on a methodology that allows us to show that the lysine-deficient proteins can undergo non-lysine ubiquitination and proteasomal degradation independent of ubiquitination. Supporting this assumption, recent Trim-Away assay results showed that the TRIM21 E3 efficiently degrades lysine-less substrates, potentially in tandem with a non-canonical ubiquitination mechanism^73^. This suggests that UPS may recruit specialised E3s that control the abundance of lysine-deficient proteins.

The presence of disordered regions in proteins that interact with the proteasome is a necessary structural requirement for ubiquitin-independent degradation^74^. Yet, VHL and SOCS1 are virtually devoid of such elements and should therefore have other features that favour ubiquitin-independent proteasomal degradation. However, as most lysine desert proteins have a high content of disordered regions, we hypothesise that they can undergo proteasomal degradation independent of ubiquitination. We envisage that our analysis and observations provide the foundation for a deeper understanding of the origin and function of lysine deserts and the non-canonical regulation of such a proteome by the UPS.

## Limitations of the study

Our analyses at this point cannot exclude potentially additional functions of lysine-free regions in proteins, like stabilization of molecular interactions or limiting non-ubiquitin modifications of the lysine. Our data also do not rule out whether autophagy or cellular extrusion pathways relaying on extracellular vesicles could be involved in the turnover of the lysine-deficient proteome. The results of our experiments do not enable us to determine whether SOCS1 and VHL undergo non-canonical ubiquitination by the CRLs of which they are components or whether other ubiquitin ligases mediate this, nor the sites of their non-lysine ubiquitination. We also could not prove that non-lysine ubiquitination of lysine-depleted proteins has a regulatory role, i.e., competing with other post-translational modifications.

## Author contributions

N.A.S. and W.P. conceived the study. M.P. performed experiments. M.P., N.A.S., P.T., and W.P. analyzed data. N.A.S. and W.P. drafted the manuscript with input from all authors.

## Figure and scheme legends

**SFigure 1. Lysine deserts in bacteria are most prevalent in *Actinobacteria* and their phages. (A)** Phylogenetic tree of different bacteria classes with a calculated average protein fraction with lysine desert min. 150 aa in the proteomes of their member taxons. Only classes with at least 10 member taxons were considered. The color gradient corresponds to the min-max normalization where red denotes classes with the lowest average fraction of desert min. 150 aa and green with the highest. **(B)** Phylogenetic tree of selected bacteria taxons from distinct classes with a calculated fraction of proteins with lysine desert min. 150 aa in their proteomes. **(C)** Bar plot of fractions of lysine desert min. 150 aa in pan proteome of selected phages’ groups. The number of sequences with lysine desert and the total number of analyzed sequences are indicated to the right of each bar.

**SFigure 2. Lysine deserts in eukaryotes ascend with growing organismal complexity. (A)** Phylogenetic tree of selected eukaryotes taxons with a calculated fraction of proteins with lysine desert min. 150 aa in their proteomes. **(B)** Phylogenetic tree of selected taxonomic families/order, including model organisms used in the previous analysis, with calculated fractions of conserved lysine desert min. 150 aa among their OGs. **(C)** Heatmap of fractions of proteins with desert region min. 150 aa of each of 20 aa among proteomes of selected eukaryotic model organisms. **(D)** Heatmaps of relative fractions of each amino acid in sequences of selected eukaryotic model organisms with a lysine desert min. 150 aa normalized to the entire appropriate proteome; value of 1.00 indicates no change. Up - considering whole sequences; down - considering only the lysine desert regions. **(E)** Histogram of distribution of sequence location of lysine desert min. 150 aa regions in the human proteome. **(F)** Box plots of predicted disorder score of residues constituting lysine desert region min. 150 aa only vs. residues of all proteins in the human proteome. Up - sequence-based disorder predictions obtained using the IUPred3 software; higher values indicate a higher disorder probability. Down - structure-based disorder predictions based on the pLDDT values obtained for the AlphaFold2 models of the human proteome; lower values indicate a higher probability of disorder. Disorder cut-offs proposed by the method’s authors are marked with dashed lines. The stars denote the significance levels per two-tailed *p*-value obtained from the Mann-Whitney U rank test. **(G)** Scatter plot showing coverage of residues building structural lysine deserts within residues constituting sequence lysine deserts in human proteome based on the AlphaFold2 models. The Spearman rank-order correlation coefficient is denoted below the plot.

**SFigure 3. Lysine deserts often co-occur with a lysine cluster. (A)** Bar plot of the fraction of lysine clusters’ occurrence among proteins with lysine desert min. 150 aa from selected model organisms. Exact values are denoted above each bar. **(B)** Bar plot of the fraction of lysine clusters’ occurrence in proteomes of selected model organisms. Exact values are denoted above each bar. **(C)** GO molecular function terms found to be associated with proteins with lysine desert min. 150 aa (up) and with proteins with lysine desert min. 150 aa along co-occurring with a lysine cluster (down). (**D**) Box plots of predicted disorder score of residues constituting lysine desert regions min. 50% only in human RING E3s vs. residues of all human RING E3s. Up - sequence-based disorder predictions obtained using the IUPred3 software; higher values indicate a higher disorder probability. Down - structure-based disorder predictions based on the pLDDT values obtained from the AlphaFold2 models of the human proteome; lower values indicate a higher probability of disorder. Disorder cut-offs proposed by the authors of the methods are marked with dashed lines. The stars denote the significance levels per two-tailed *p*-value obtained from the Mann-Whitney U rank test. (**E**) Box plots of predicted disorder score of residues constituting lysine desert region min. 150 aa only in human RING E3s vs. residues of all in human RING E3s. Up - sequence-based disorder predictions obtained using the IUPred3 software; higher values indicate a higher disorder probability. Down - structure-based disorder predictions based on the pLDDT values obtained for the AlphaFold2 models of the human proteome; lower values indicate a higher probability of disorder. Disorder cut-offs proposed by the method’s authors are marked with dashed lines. The stars denote the significance levels per two-tailed *p*-value obtained from the Mann-Whitney U rank test. **(F)** Histogram of distribution of sequence location of lysines within proteins with lysine desert min. 150 aa from the human proteome.

**SFigure 4. 20 µM concentration of MG132 proteasome inhibitor does not cause cytotoxic effects.** Cell viability during the NanoBret ubiquitination assays was evaluated using the CellTiter-Glo 2.0 Cell Viability Assay (for details, see Materials and Methods). **(A)** Treatment of 20 µM MG132 of VHL and its lysine-less variant. **(B)** Treatment of 20 µM MG132 of SOCS1 and its lysine-less variant. The significance levels obtained from Šídák’s multiple comparisons test were indicated for the compared conditions (ns - not significant). Analyzed and visualized in the GraphPad Prism 9.

**SFigure 5. Lysine deserts do not correlate with protein half-life.** Scatter plots showing a lack of correlation between the length of the lysine-less region expressed as nominal value and protein half-life in eight different cell types from two large-scale proteomic studies of human proteins. The Spearman rank-order correlation coefficient is denoted below each plot.

**Table S1.** Proteomes from the UniProt database used in the lysine desert analyses.

**Table S2.** Summary of pupylomes of M. tuberculosis, M. smegmatis, and C. glutamicum.

**Table S3.** Summary of occurrence of lysine desert min. 50% in pupylomes of *M. tuberculosis*, *M. smegmatis,* and *C. glutamicum*.

**Table S4.** Summary of occurrence of lysine desert min. 150 aa in pupylomes of *M. tuberculosis*, *M. smegmatis,* and *C. glutamicum*.

**Table S5.** Summary of occurrence of lysine desert min. 50% in non-pupylated proteins of *M. tuberculosis*, *M. smegmatis,* and *C. glutamicum*.

**Table S5.** Summary of occurrence of lysine desert min. 150 aa in non-pupylated proteins of *M. tuberculosis*, *M. smegmatis,* and *C. glutamicum*.

**Table S7.** Human proteome with calculated lysine deserts, lysine clusters, and other relevant information.

**Table S8.** Three most extended structural lysine deserts calculated for the human proteome. **Table S9.** Orthologous groups from *Saccharomycetaceae* with conserved lysine desert min. 50%.

**Table S10.** OGs from *Drosophilidae*, *Rodentia*, and *Hominidae* (in *Saccharomycetaceae* and *Rhabditida*, OGs were either absent or, if present, conserved lysine desert min. 50% also needed to occur) with conserved lysine desert min. 50%.

**Table S11.** Human E3 ligases with their annotated types.

**Table S12.** Orthologs of human E3 ligases with ≤ 5 lysines in *D. rerio*, *X. silurana*, *G. gallus* and *M. musculus*.

**Table S13.** Relative solvent accessibility (RSA) values for lysines of human E3 ligases with ≤ 5 lysines calculated using the AlphaFold2 models.

## Materials and Methods

### Analysis of lysine deserts in bacteria and eukaryotic proteomes

All bacteria and eukaryotic reference proteomes (for 8881 and 1329 taxons, respectively) were downloaded from the UniProt database^40^ (from the FTP repository; data obtained on 25.05.2022; see Table S1 for the summary of downloaded data). In all performed analyses, sequences <150 aa were excluded. For selected taxons, namely, *M. tuberculosis* H37Rv (virulent), *M. smegmatis*, *C. glutamicum*, *S. ceolicolor*, *L. ferrooxidans*, *B. subtilis*, *E. coli*, *S. cerevisiae*, *C. elegans*, *D. melanogaster*, *M. musculus*, and *H. sapiens*, number of transmembrane helices (TMH) for each sequence in their proteome were predicted using TMHMM-2.0 software^41, 42^. For analyses concerning these taxons, proteins with a predicted number of TMH >2 were excluded. Summary of a number of sequences prior and after filtering can be found in Table 1 and Table 3. Sequences with a lysine desert of a declared type (lysine-less region of min. 150 aa or constituting min. 50% of the sequence) were counted in each proteome and a fraction of sequences with a given lysine desert was reported for each taxon or averaged for proteomes of the same bacteria class (only classes with at least 10 taxons were considered).

### Analysis of bacteria pupylomes

The dataset of pupylated proteins of *M. tuberculosis*, *M. smegmatis,* and *C. glutamicum* was downloaded from the PupDB database^44^ (data obtained on 04.08.2022). As some of the UniProt IDs in the obtained dataset were obsolete and could not be mapped directly to the UniProt reference proteomes of selected taxons, the UniProt retrieve the correct UniProt IDs. Among pupylated and non-pupylated proteins of the aforementioned taxons, sequences <150 aa and with >2 predicted TMH were excluded from further analyses (all proteins from the UniProt reference proteome of given taxon were considered as non-pupylated except those reported in the PupDB). The remaining proteins were screened for the presence of a lysine desert.

### Analysis of lysine deserts in phages’ proteomes

Proteomes of Mycobacterium, Escherichia and Bacillus phages were downloaded from the UniProt database using the mycobacterium AND (taxonomy_id:10239), escherichia AND (taxonomy_id:10239), and bacillus AND (taxonomy_id:10239) queries, respectively (data obtained on 31.07.2022). Proteomes marked as outliers as well as those with <40 sequences were excluded from further analyses (see the summary of proteomes selected for analysis in Table S1). Next, for each phages’ group separately, all sequences, excluding those <150 aa, were concatenated into one fasta file and clustered using the cd-hit web server^75^ (with the default parameters, identity cutoff =0.9) to create non-redundant pan proteome (summary of pan proteomes’ properties is presented in Table 2). Sequences with a lysine desert of declared type were counted in each pan proteome and a fraction of sequences with a given lysine desert was reported.

### Lysine desert search within the Orthologous Groups

#### Dataset

Lysine desert search among the Orthologous Groups (OGs) of *Saccharomycetaceae*, *Rhabditida*, *Drosophilidae*, *Rodentia*, and *Hominidae* was performed based on the datasets from the eggNOG5 database^45^ (data obtained on 31.07.2022).

#### Data preprocessing

For each mentioned taxonomic family/order, we selected Multiple Sequence Alignment (MSA) files of their OGs, which fulfilled the following conditions: (i) the presence of minimum four sequences, (ii) the presence of a minimum of 60% of the taxonomic family’s/order’s taxons (each taxonomic family/order from the eggNOG5 database owned a maximum number of taxons for which MSA files were constructed, however not all MSA files cover all taxons), (iii) median protein length of min. 150 aa, (iv) mean a number of predicted TMHs using the TMHMM-2.0 software ≤ 2, (v) the presence of at least one sequence from the representative organism for the particular taxonomic family/order - representative organisms are: *S. cerevisiae*, *C. elegans*, *D. melanogaster*, *M. musculus*, and *H. sapiens*. This data filtering resulted in the analysis of 50-68% of all available OGs (Table 4).

#### Definition of a conserved lysine desert

For the purpose of OG-based analyses, we broadened the definition of the lysine desert, so the OG is considered as containing a conserved lysine desert if (i) at least 60% of its sequences possess a lysine desert region of a minimum length of 150 aa, or (ii) at least 60% of its sequences possess a lysine desert region constituting a minimum of 50% of OG’s median sequence length.

#### Finding analogous OGs between different families/orders

The eggNOG5 database does not provide direct information on which OGs of different taxonomic ranks cover analogous proteins. Therefore, such “vectors” of corresponding MSAs from analogous OGs were derived based on *Opisthokonta* - a broad eukaryotic supergroup, including animals and fungi. Briefly, over 100.000 MSAs of OGs belonging to *Opisthokonta*, also obtained from the eggNOG5 database, were screened to find sequences of analogous proteins from *S. cerevisiae*, *C. elegans*, *D. melanogaster*, *M. musculus*, and *H. sapiens* and match them with their presence in OGs of *Saccharomycetaceae*, *Rhabditida*, *Drosophilidae*, *Rodentia*, and *Hominidae.* The analyses were not performed directly on the OGs from *Opisthokonta*, as although their MSAs cover all aforementioned model organisms, they also usually contain long gaps regions (due to coverage of multiple evolutionary distant organisms) which would not allow for comparison lysine desert conservation between selected families/orders. The IDs of corresponding MSAs of OGs of *Saccharomycetaceae*, *Rhabditida*, *Drosophilidae*, *Rodentia*, and *Hominidae* are available at https://github.com/n-szulc/lysine_deserts.

### Disorder predictions

The probability of each residue being disordered was calculated for all sequences in the human proteome, excluding those <150 aa and with the predicted number of TMH >2. Disorder predictions were obtained using two approaches: (i) by running the standalone version of the IUPred3^47^ software with the default parameters, (ii) by obtaining the pLDDT values from the AlphaFold2 models^49^ of the human proteome downloaded from the AlphaFold Protein Structure Database^50^ (proteome ID: UP000005640_9606_HUMAN_v3; since AlphaFold2 models of proteins >2700 aa are split to separate overlapping files, such proteins were also excluded from the analysis).

### Structural lysine desert algorithm

#### Dataset

The structural lysine desert screen was performed on human proteome downloaded from the AlphaFold Protein Structure Database (proteome ID: UP000005640_9606_HUMAN_v3; since AlphaFold2 models of proteins >2700 aa are split to separate files, such proteins were also excluded from the analysis).

#### Data preprocessing

Each protein structure was parsed using the Biopython^76^ python module (v. 1.79) and list of contacts (neighbors) (default contact cut-off =4.0 Å; defined as the shortest euclidean distance between any heavy atoms of the two residues) together with solvent-accessible surface area (SASA) and relative solvent accessibility (RSA) values obtained using the DSSP program^77^ (v. 3.0.0); Sander method^78^ was used for SASA normalization to acquire RSA) were calculated for each residue.

#### Aim

The algorithm intends to find the longest, uninterrupted lysine-less regions among solvent-exposed residues that remain in contact. Therefore, buried residues may also break the continuity of the structural lysine desert, as there may be no exposed neighbors to spread to (cases visible in Fig 3G when the entire protein is lysine-less yet the structural lysine desert does not equal 100%).

#### Algorithm

The complete code and documentation, including the algorithm’s visualization, are available at https://github.com/n-szulc/lysine_deserts. Briefly, all contacts are analyzed for each residue that is not a lysine and is solvent-exposed (RSA cut-off value >0.2). If there is no solvent-exposed lysine among these contacts, the exposed contacts are added to the temporary list containing residues of the structural lysine desert; otherwise, the algorithm stops. Next, all contacts of the aforementioned contacts are analyzed. If there is no solvent-exposed lysine among them, the exposed ones are added to the temporary list containing residues of the structural lysine desert. The algorithm stops if such exposed ones are absent or a solvent-exposed lysine occurs. The algorithm repeats and saves the three most extended structural lysine deserts.

#### Remarks

When iterating over contacts, if a residue has no calculated SASA or RSA value due to the DSSP error, it is omitted, and the algorithm proceeds to the next one, except for lysine. In such a case, lysine is always considered solvent-exposed.

### Analysis of the correlation of length of lysine desert and protein half-life

The protein half-lives datasets were obtained from recent high-throughput proteomic studies^56, 57^. The dataset from Mathieson and colleagues provided information on the turnover of human proteins in primary cells - B cells, NK cells, hepatocytes, and monocytes, while the dataset from Li and colleagues outlined half-lives of short-lived human proteins measured in four cell lines - U2OS, HEK293T, HCT116, and RPE1. Mathieson and colleagues usually reported half-live values from two replicates for a given type of primary cells, mean protein half-life was calculated in all such cases; Li and colleagues reported only one value of the protein half-life in a given cell line. Gene symbols reported by Mathieson and colleagues and UniProt IDs provided by Li and colleagues were mapped to the UniProt reference human proteome to associate half-life values with representative proteins; half-life values whose identifier was unmapped were discarded from further analyses. Longest lysine-less region was calculated in all human proteins, excluding those <150 aa, with predicted (using the TMHMM-2.0 software) number of TMH >2, and with outlier half-life values (from 0.01 and 0.99 quantiles), and expressed as either a nominal value or sequence fraction. A summary of the number of analyzed proteins with assigned half-life values is presented in Table 5.

**Table 5.** Summary of the number of analyzed proteins with a measured half-life in each dataset.

|  | Number of proteins | Number of proteins mapped to the reference human proteome | Number of proteins used in the analysis<sup>a</sup> |
| --- | --- | --- | --- |
| **Mathieson *et al.*, 2018** |  |  |  |
| B cells | 4653 | 4134 | 3885 |
| NK cells | 3555 | 3203 | 3005 |
| Hepatocytes | 4242 | 3755 | 3342 |
| Monocytes | 4649 | 4193 | 3886 |
| **Li *et al.*, 2021** |  |  |  |
| U2OS | 1428 | 1175 | 1140 |
| HEK293T | 1434 | 1151 | 1124 |
| HCT116 | 1904 | 1571 | 1520 |
| RPE1 | 1652 | 1335 | 1278 |

<sup>a</sup> Excluding sequences <150 aa, with the predicted number of TMH >2, and with outlier half-life values (from 0.01 and 0.99 quantile).

### Over-representation analysis

The GO-based over-representation analyses of human lysine desert min. 150 aa/min. 50% proteins, regardless of and along with lysine clusters, were performed using the WebGestalt web server with default parameters^79^; our filtered human proteome served as background. The false discovery rate (FDR) was controlled to 0.05 using the Benjamini-Hochberg method for multiple testing. The results were visualized as treemaps using the modified R scripts generated by the REVIGO web server^80^ (species specified as *Homo sapiens*, the rest of parameters was set as default).

### Plasmid construction

The sequence and ligation independent cloning (SLIC) method^81^, was used to construct HiBiT and NanoLuc fusion vectors. HiBiT fusion vectors were constructed using the parental vector pBiT3.1-N (N2361, Promega). To generate the HiBiT-VHL vector, the pBiT3.1-N vector was linearized with SacI, and HindIII enzymes, dephosphorylated and mixed with human VHL sequence PCR-amplified from HEK293 cDNA. Synthesis of HiBiT-SOCS1 was ordered from Azenta Life Sciences. To obtain HiBiT-SOCS1 K118R and HiBiT-VHL K159R, K171R, and K196R variants, splice-PCRs were performed using HiBiT-SOCS1 and HiBiT-VHL vectors as templates, respectively. NanoLuc fusion vectors were constructed using the parental vectors pNLF1-N (N1351, Promega) and pNLF1-C (N1361, Promega) to obtain N-terminal and C-terminal NanoLuc-tagged protein fusions, respectively. To generate NanoLuc-VHL and NanoLuc-SOCS1 vectors, the pNLF1-N vectors were linearized with XbaI and XhoI enzymes, dephosphorylated, and linearized vectors were mixed with VHL or SOCS1 sequences, which were PCR-amplified using HiBiT-VHL or HiBiT-SOCS1 vectors as templates. For VHL and SOCS1 cloning into pNLF1-C vectors, the vector was linearized with PvuI and XhoI enzymes. In order to construct NanoLuc-tagged VHL K159R, K171R, K196R, and SOCS1 K118R mutated variants, SLIC cloning was performed using VHL and SOCS1 sequences, which were PCR-amplified using HiBiT-VHL (K159R, K171R, K196R) and HiBiT-SOCS1 (K118R) vectors, respectively, and the linearized pNLF1-N and pNLF1-C vectors. The list of primers’ sequences used to generate the constructs is available in Table 6.

**Table 6.**
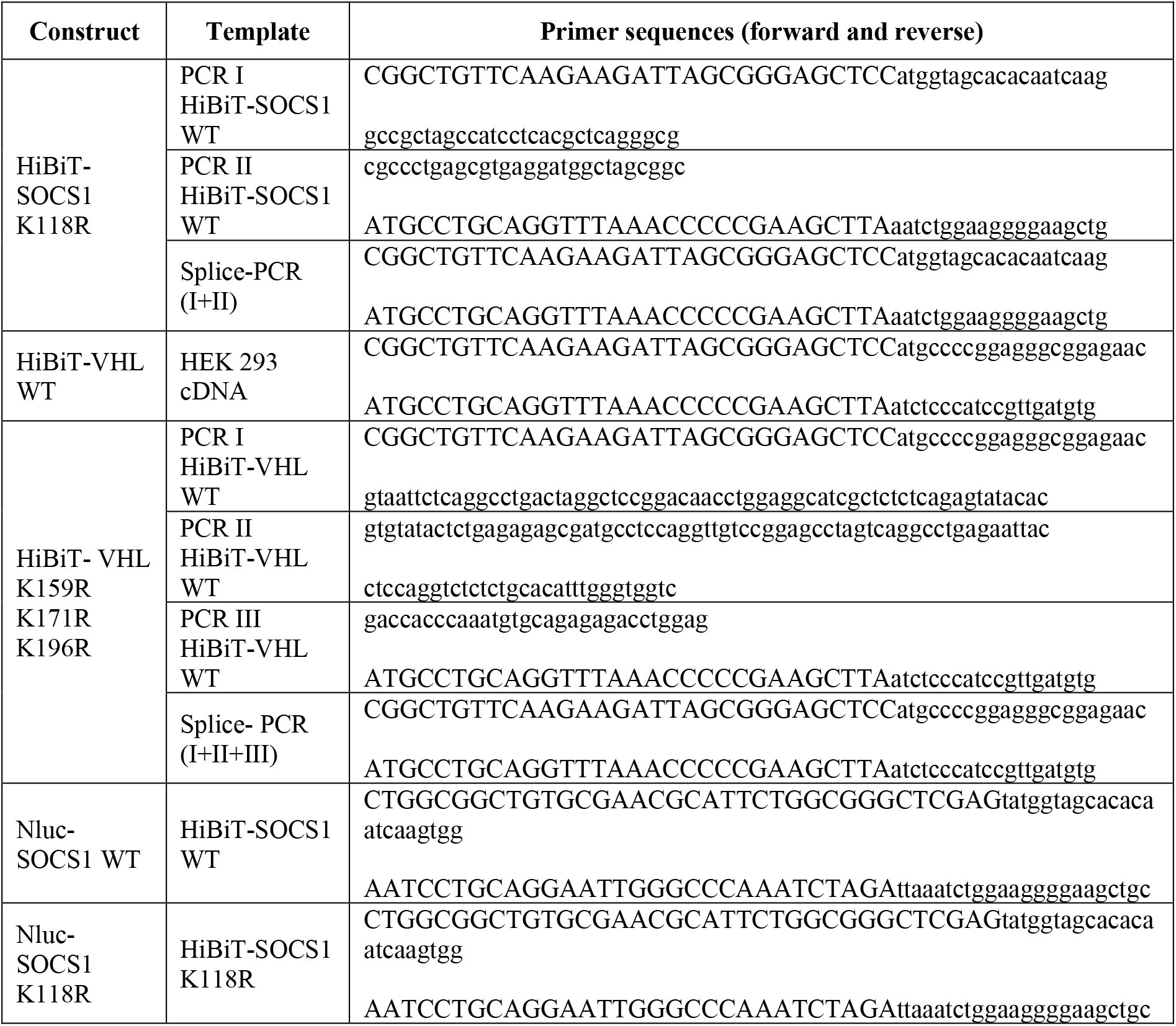
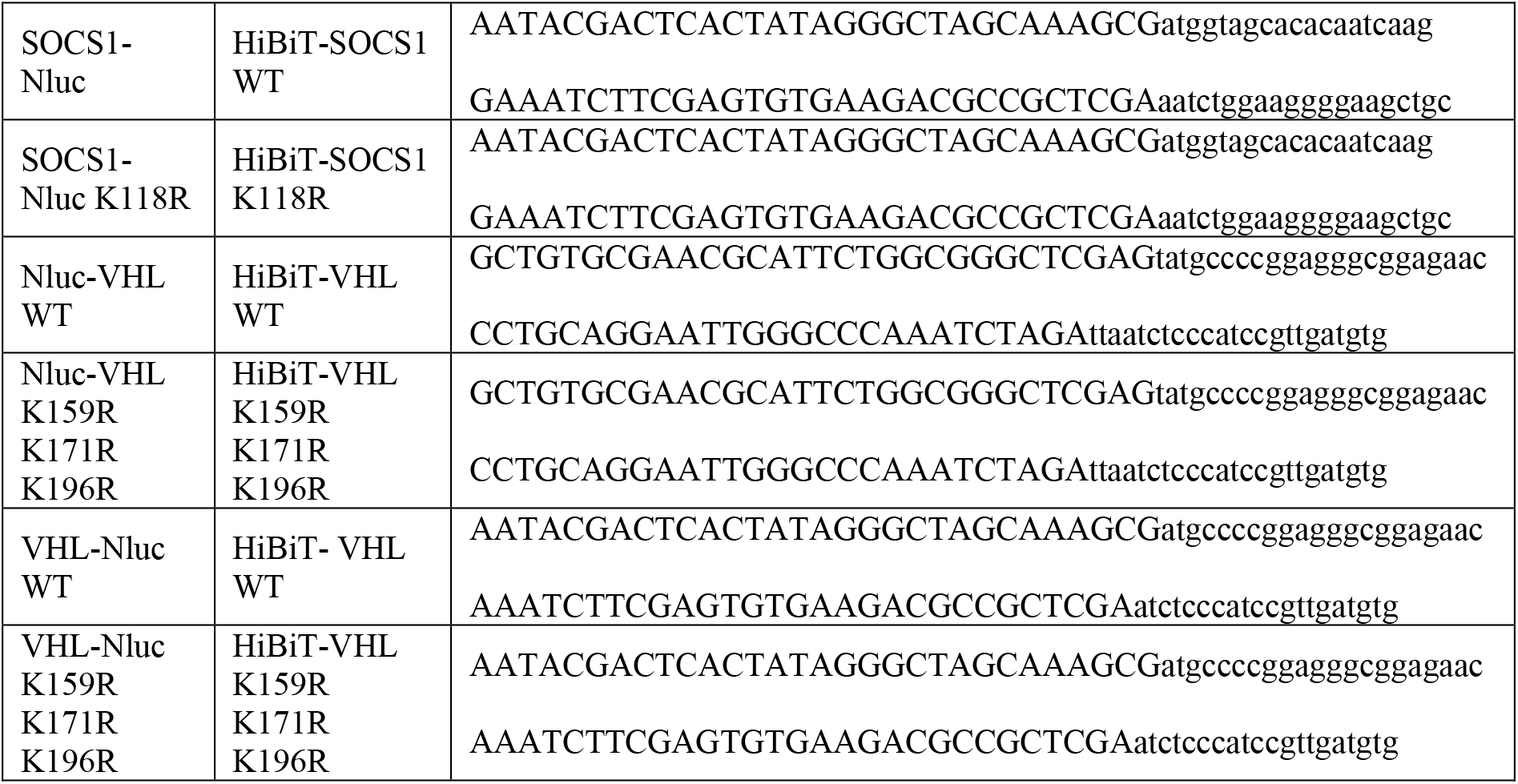
List of primers used in this study for cloning.

### Cell culture

Flp-In 293 cell line (HEK293; ThermoFisher Scientific, product no. 510021, lot no. 2348919) were cultivated in Dulbecco’s Modified Eagle’s Medium (DMEM; D6429, Sigma) supplemented with 10% heat-inactivated Fetal Bovine Serum (FBS; F9665, Sigma) and 1% antimycotic antibiotic (15240062, Gibco) at 37°C, 5% CO_2_ in a humidified incubator.

### NanoBRET ubiquitination assay

#### Cells preparation

HEK293 cells were seeded in 6-well plates at 800.000 cells per well. After 6-8 hours, cells were transiently transfected with 1 or 2 µg HaloTag-Ubiquitin (N2721, Promega) or the control vector pHTN HaloTag CMV-neo (G7721, Promega) and 0.01 or 0.02 µg NanoLuc-tagged SOCS1 or VHL expression constructs, respectively (acceptor to donor ratio was maintained at the ratio of 100:1). Transfection was carried out using the FuGENE HD Transfection Reagent (E2312, Promega) according to the manufacturer’s guidelines, in the Opti-MEM I Reduced Serum Medium, no phenol red (11058021, Gibco), maintaining the 3:1 FuGENE HD:DNA ratio. After 20-22 hours, cells were trypsinized, counted, and resuspended in the Opti-MEM containing 4% FBS and 1% antimycotic antibiotic at a concentration of 200.000 cells/ml.

#### NanoBret assay

The bioluminescence resonance energy transfer (BRET) ubiquitination assay was performed using the NanoBRET Nano-Glo Detection Systems (Standard) (N1661, Promega) according to the manufacturer’s guidelines. Briefly, the HaloTag 618 ligand was added to the cells (at 100 nM concentration), and 100 μl of the cell suspension was plated per well of white 96-well tissue culture plates (655083, Greiner). Dimethyl sulfoxide (DMSO) was added (1 μl/ml) instead of the HaloTag 618 ligand to the control cells. After 20-22 hours, cells were treated with 20 µM MG132 (S2619, SelleckChem) or DMSO (for vehicle control) for 2 hours. For the compound preparation, 40 mM of the MG132 stock solution in DMSO was diluted in the Opti-MEM to 400 µM concentration (20x), and 5 μl of the solution was added to appropriate wells. After the treatment, the NanoBRET Nano-Glo Substrate diluted in Opti-MEM was added to wells, and donor and acceptor light emissions were immediately measured using the TECAN Infinite 200 Pro plate reader equipped with the Magellan Pro software with the Blue2 NB and the Red NB filters, and integration time of 1000 ms. BRET ratios in mBRET units were calculated by dividing the acceptor emission (Red NB) by the donor luminescence (Blue2 NB) and multiplied by 1000. The values were corrected by subtracting the BRET ratio measured in equivalent samples which received DMSO instead of the HaloTag 618 ligand.

#### Cell viability assay

To assess cell viability during the NanoBret assays, they were multiplexed with the CellTiter-Glo 2.0 Cell Viability Assay (G9241, Promega) following the manufacturer’s guidelines. Briefly, an equal volume (125 μl) of the CellTiter-Glo 2.0 Reagent was added to each well and mixed on a plate shaker for 5 min. After a 30-min incubation, luminescence was measured using the TECAN Infinite 200 Pro plate reader equipped with the Magellan Pro software and an integration time of 1000 ms. The untransfected cells were used as the global viability reference.

### Cycloheximide chase assay

#### Cells preparation

HEK293 cells were seeded in white 96-well tissue culture plates (655083, Greiner) at a density of 10.000 cells in a total volume of 100 µl per well. After 38-40 hours, cells were transiently transfected with 2.5 ng HiBiT-tagged VHL or SOCS1 expression constructs, diluted in carrier DNA (E4881, Promega) to obtain a final DNA amount of 50 ng/well. Transfection was carried out using the FuGENE HD Transfection Reagent (E2312, Promega) according to the manufacturer’s guidelines in the Opti-MEM I Reduced-Serum Medium (31985047, Gibco), maintaining the 3:1 FuGENE HD: DNA ratio. Cells were incubated for 23 hours at 37°C, 5% CO_2_.

#### Cycloheximide chase assay

Cycloheximide (CHX; CYC003.5, BioShop) was added to wells at a final 50 µg/ml concentration for up to 4 hours. Treatment was performed in triplicate. Where indicated, cells were concomitantly treated with 20 µM MG132 (S2619, SelleckChem) for 4 hours and 1 µM TAK-243 (S8341, SelleckChem) for 5 hours. TAK-243 was added to wells 1 hour before the first dose of CHX. After a 4-hour incubation with CHX, the Nano-Glo HiBiT Lytic Detection Assay (N3040, Promega) was performed according to the manufacturer’s guidelines. Briefly, 120 µl of the Nano-Glo HiBiT Lytic Reagent was added to each well. Samples were mixed on an orbital shaker for 5 min and incubated for 10 min before luminescence measurement using the TECAN Infinite 200 Pro plate reader equipped with the Magellan Pro software and an integration time of 1000 ms. The luminescence measurements were normalized in each well to the number of living cells (see below).

#### Cell viability assay

To assess cell viability during the CHX assays, they were multiplexed with the CellTiter-Fluor Cell Viability Assay (G6080, Promega) following the manufacturer’s guidelines. Briefly, after 3-hour incubation with CHX, 20 µl of the 5x concentrated CellTiter-Fluor Reagent was added to all wells. Cells were incubated at 37°C for 1 hour, and fluorescence was measured using the Tecan Infinity M1000 fluorescence plate reader equipped with the Magellan Pro software with the parameters setup of 390nm_Ex_/505nm_Em_. The untransfected cells were used as the global viability reference.

### Key resources table

| REAGENT or RESOURCE | SOURCE | IDENTIFIER |
| --- | --- | --- |
| Chemicals, peptides, and recombinant proteins |  |  |
| Cycloheximide | BioShop | Cat#CYC003.5 |
| MG132 | SelleckChem | Cat#S2619 |
| TAK-243 | SelleckChem | Cat#S8341 |
| FuGENE HD Transfection Reagent | Promega | Cat#E2312 |
| Critical commercial assays |  |  |
| Nano-Glo HiBiT Lytic Detection Assay | Promega | Cat#N3040 |
| CellTiter-Fluor Cell Viability Assay | Promega | Cat#G6080 |
| NanoBRET Nano-Glo Detection Systems | Promega | Cat#N1661 |
| CellTiter-Glo 2.0 Cell Viability Assay | Promega | Cat#G9241 |
| Deposited data |  |  |
| https://github.com/n-szulc/lysine_deserts<sup>52</sup> | This work | N/A |
| Experimental models: Cell lines |  |  |
| Human: Flp-In 293 | ThermoFisher Scientific | Cat#R75007 |
| Oligonucleotides |  |  |
| Primers for VHL and SOCS1 cloning and mutagenesis, see Table 6 | This work | N/A |
| Recombinant DNA |  |  |
| HiBiT-SOCS1 WT | Azenta Life Sciences | N/A |
| HiBiT- and NanoLuc-tagged constructs, see Table 6 | This work | N/A |
| HaloTag-Ubiquitin | Promega | Cat#N2721 |
| pHTN HaloTag CMV-neo | Promega | Cat#G7721 |
| pBiT3.1-N | Promega | Cat#N2361 |
| pNLF1-N | Promega | Cat#N1351 |
| pNLF1-C | Promega | Cat#N1361 |
| Software and algorithms |  |  |
| cd-hit web server | http://cd-hit.org | N/A |
| DSSP (v. 3.0.0) used within the Biopython python module | https://anaconda.org/salilab/dssp | N/A |
| GraphPad Prism 9 (v. 9.5.0) | https://www.graphpad.com | N/A |
| IUPred3 standalone | https://iupred3.elte.hu | N/A |
| Magellan Pro | https://lifesciences.tecan.com/software-magellan | N/A |
| Own codes | https://github.com/n-szulc/lysine_deserts | N/A |
| REVIGO web server | http://revigo.irb.hr | N/A |
| TMHMM-2.0 standalone | https://services.healthtech.dtu.dk/service.php?TMHMM-2.0 | N/A |
| UniProt Retrieve/ID mapping | https://www.uniprot.org/id-mapping | N/A |
| WebGestalt web server | http://www.webgestalt.org | N/A |

### Data analysis and plotting

Statistical tests and correlation coefficients regarding the bioinformatics analyses were calculated using the SciPy^82^ (v. 1.7.3) python module. Phylogenetic trees were plotted using the ETE python toolkit^83^ (v. 3.1.2). Results of the CHX and NanoBret assays were analyzed and visualized in the GraphPad Prism 9 (v. 9.5.0). All other plots were generated using the matplotlib (v. 3.5.1) and seaborn (v. 0.11.2) python modules. Graphics were created with BioRender.com.

## Data availability

The code required for performing all the described analyses, datasets generated during this study, and raw luminescence and fluorescence measurements from the CHX and NanoBret assays can be found in the repository at https://github.com/n-szulc/lysine_deserts52.

## Acknowledgments

We thank the Genome Engineering Unit of the International Institute of Molecular and Cell Biology in Warsaw (geu.iimcb.gov.pl) for generating DNA constructs. We would also like to sincerely thank Marta Niklewicz for her technical assistance. We thank Dr. Filip Stefaniak for sharing his computational resources. We acknowledge Dr. Jan Ludwiczak for his initial bioinformatic analysis. We warmly thank Prof. Dario Valenzano from the Leibniz Institute on Aging (FLI) and Dr. Milka Kostic from the Dana-Farber Cancer Institute for their insightful comments on the study. For the purpose of Open Access, the authors have applied a CC-BY public copyright licence to any Author Accepted Manuscript (AAM) version arising from this submission.

