## Supplementary Tables 1-13 for "Lysine-deficient proteome can be regulated through non-canonical ubiquitination and ubiquitin-independent proteasomal degradation": Supplementary Information.pdf

**for**

**Table S2.** Summary of pupylomes of *M. tuberculosis*, *M. smegmatis* and *C. glutamicum*.

| | Number of proteins in proteome | Number of pupylated proteins | Fraction of pupylated proteins |
|---|---|---|---|
| ***M. tuberculosis*** | 3995 | 54 | 1.35 |
| ***M. smegmatis*** | 6602 | 76 | 1.15 |
| ***C. glutamicum*** | 3093 | 43 | 1.39 |

**Table S3.** Summary of occurrence of lysine desert min. 50% in pupylomes of *M. tuberculosis*, *M. smegmatis* and *C. glutamicum*.

| | Total number of sequences | Total number of filtered[a] sequences | Average length of filtered[a] sequences [aa] | Number of sequences with lysine desert min. 50% | Fraction of sequences with lysine desert min. 50% |
|---|---|---|---|---|---|
| ***M. tuberculosis*** | 54 | 49 | 405.86 | 4 | 8.16 |
| ***M. smegmatis*** | 76 | 66 | 406.83 | 1 | 1.52 |
| ***C. glutamicum*** | 43 | 39 | 370.23 | 1 | 2.56 |

[a] Excluding sequences <150 aa and with the predicted number of TMH >2.

**Table S4.** Summary of occurrence of lysine desert min. 150 aa in pupylomes of *M. tuberculosis*, *M. smegmatis* and *C. glutamicum*.

| | **Total number of sequences** | **Total number of filtered[a] sequences** | **Average length of filtered[a] sequences [aa]** | **Number of sequences with lysine desert min. 150 aa** | **Fraction of sequences with lysine desert min. 150 aa** |
|---|---|---|---|---|---|
| ***M. tuberculosis*** | 54 | 49 | 405.86 | 8 | 16.33 |
| ***M. smegmatis*** | 76 | 66 | 406.83 | 2 | 3.03 |
| ***C. glutamicum*** | 43 | 39 | 370.23 | 1 | 2.56 |

[a] Excluding sequences <150 aa and with the predicted number of TMH >2.

**Table S5.** Summary of occurrence of lysine desert min. 50% in non-pupylated proteins of *M. tuberculosis*, *M. smegmatis* and *C. glutamicum*.

| | **Total number of sequences** | **Total number of filtered[a] sequences** | **Average length of filtered[a] sequences [aa]** | **Number of sequences with lysine desert min. 50%** | **Percentage of sequences with lysine desert min. 50%** |
|---|---|---|---|---|---|
| ***M. tuberculosis*** | 3940 | 2729 | 386.62 | 710 | 26.02 |
| ***M. smegmatis*** | 6524 | 4601 | 355.82 | 1223 | 26.58 |
| ***C. glutamicum*** | 3050 | 1948 | 360.01 | 185 | 9.5 |

[a] Excluding sequences <150 aa and with the predicted number of TMH >2.

**Table S6.** Summary of occurrence of lysine desert min. 150 aa in non-pupylated proteins of *M. tuberculosis*, *M. smegmatis* and *C. glutamicum*.

| | Total number of sequences | Total number of filtered[a] sequences | Average length of filtered[a] sequences [aa] | Number of sequences with lysine desert min. 150 aa | Percentage of sequences with lysine desert min. 150 aa |
|---|---|---|---|---|---|
| ***M. tuberculosis*** | 3940 | 2729 | 386.62 | 842 | 30.85 |
| ***M. smegmatis*** | 6524 | 4601 | 355.82 | 1341 | 29.15 |
| ***C. glutamicum*** | 3050 | 1948 | 360.01 | 157 | 8.06 |

[a] Excluding sequences <150 aa and with the predicted number of TMH >2.

**Table S12.** Orthologs of human E3 ligases with ≤ 5 lysines[a] in *D. rerio*, *X. silurana*, *G. gallus* and *M. musculus*.

| D. rerio E3 ligase | D. rerio UniProt ID | D. rerio no of lysines | X. silurana E3 ligase | X. silurana UniProt ID | X. silurana no of lysines | G. gallus E3 ligase | G. gallus UniProt ID | G. gallus no of lysines | M. musculus E3 ligase | M. musculus UniProt ID | M. musculus no of lysines | H. sapiens E3 ligase | H. sapiens UniProt ID | H. sapiens no of lysines | CRL receptor |
|---|---|---|---|---|---|---|---|---|---|---|---|---|---|---|---|
| Kbtbd13a; Kbtbd13b | A0A140LGF5; X1WCK8 | 28; 20 | Kbtbd13 | F6U0R1 | 22 | KBTBD13 | A0A8V0Z2K6 | 14 | Kbtbd13 | Q8C828 | 5 | KBTBD13 | C9JR72 | 4 | yes |
| Socs1A; Socs1B | Q6DEF9;E7FDH8 | 15;11 | Socs1 | Q5M8D9 | 15 | SOCS1 | B6RCQ2 | 13 | Socs1 | O35716 | 1 | SOCS1 | O15524 | 1 | yes |
| Vhl; Vhll | A1L296;B3DHE4 | 5;7 | Vhl | A9ULN6 | 6 | VHL | A0A8V0Z8B2 | 6 | Vhl | P40338 | 4 | VHL | P40337 | 3 | yes |
| - | - | - | - | - | - | - | - | - | Kctd11 | Q8K485 | 2 | KCTD1 | Q693B1 | 1 | yes |

| | | | | | | | | | | | | | | | |
|---|---|---|---|---|---|---|---|---|---|---|---|---|---|---|---|
| | | | | | | | | | | | | 1[b] | | | |
| Rnf126 | A2RV40 | 6 | Rnf126 | Q6DIP3 | 3 | RNF126 | A0A1L1RZV7 | 4 | Rnf126 | Q91YL2 | 4 | RNF126 | Q9BV68 | 4 | no |
| Fbxl15 | H9KUW9 | 8 | Fbxl15 | Q5XGC0 | 16 | FBXL15 | F1NF36 | 11 | Fbxl15 | Q91W61 | 2 | FBXL15 | Q9H469 | 2 | yes |
| Rnf181 | Q7ZW78 | 8 | Rnf181 | Q5M974 | 8 | - | - | - | Rnf181 | Q9CY62 | 7 | RNF181 | Q9P0P0 | 5 | no |
| Rnf6 | F1R4P2 | 3 | - | - | - | RNF6 | A0A1D5NWA7 | 4 | Rnf6 | Q9DBU5 | 4 | RNF6 | Q9Y252 | 4 | no |
| Rnf11a; Rnf11b | B8A662;B0V2S5 | 3;4 | Rnf11 | Q28H59 | 4 | RNF11 | F1NLF7 | 4 | Rnf11 | Q9QYK7 | 4 | RNF11 | Q9Y3C5 | 4 | no |
| Rnf44 | Q08CG8 | 5 | Rnf44 | A0A8J0PJJ6 | 6 | RNF44 | R4GIU9 | 5 | Rnf44 | Q8BI21 | 6 | RNF44 | Q7L0R7 | 5 | no |
| - | - | - | Siah3 (Xenbase) | A0A6I8RL86 | 9 | SIAH3 | R4GKE1 | 7 | Siah3 | B2RWG3 | 5 | SIAH3 | Q8IW03 | 5 | no |
| - | - | - | Fbxo27 (Xenbase) | Q0VA18 | 22 | - | - | - | Fbxo27 | Q6DIA9 | 8 | FBXO27 | Q8NI29 | 5 | yes |
| Neurl2 | A4IG40 | 16 | Neurl2 | F7CZK9 | 13 | NEURL2 | F1NIQ8 | 5 | Neurl2 | Q9D0S4 | 6 | NEURL2 | Q9BR09 | 5 | yes |
| Rnf224 | A5PMH4 | 11 | - | - | - | - | - | - | Rnf224 | Q3UIW8 | 3 | RNF224 | P0DH78 | 2 | no |

[a] Searched in human proteome, excluding sequences <150 aa, with the predicted number of TMH >2, or annotated as membrane-bound.

[b] Although there are no known orthologs of KCTD11 from *D. rerio*, *X. silurana*, and *G. gallus*, we also consider it as decreasing its lysine content in the evolution, as its ortholog from *C. picta* turtle has 4 lysines (UniProt ID: A0A8C3H7N5).

**Table S13.** Relative solvent accessibility (RSA) values for lysines of human E3 ligases with ≤ 5 lysines[a] calculated using the AlphaFold2 models.

| **UniProtID** | **Gene symbol** | **AlphaFold2 model ID** | **Lysines indices** | **Lysines RSA** |
|---|---|---|---|---|

| | | | | |
|---|---|---|---|---|
| Q9H469 | FBXL15 | AF-Q9H469-F1-model_v3 | 180;268 | 0.63;0.76 |
| Q8NI29 | FBXO27 | AF-Q8NI29-F1-model_v3 | 126;163;164;218;245 | 0.54;0.0;0.29;0.68;0.65 |
| C9JR72 | KBTBD13 | AF-C9JR72-F1-model_v3 | 210;222;390;427 | 0.32;0.36;1.0;0.16 |
| Q693B1 | KCTD11 | AF-Q693B1-F1-model_v3 | 31 | 0.79 |
| Q9BR09 | NEURL2 | AF-Q9BR09-F1-model_v3 | 73;237;276;279;283 | 0.3;0.19;0.69;0.2;0.14 |
| Q9Y3C5 | RNF11 | AF-Q9Y3C5-F1-model_v3 | 6;82;94;95 | 1.0;0.73;0.98;0.6 |
| Q9BV68 | RNF126 | AF-Q9BV68-F1-model_v3 | 207;209;233;271 | 0.55;0.66;0.74;0.57 |
| Q9P0P0 | RNF181 | AF-Q9P0P0-F1-model_v3 | 56;75;109;134;137 | 0.76;0.58;0.77;0.58;0.64 |
| P0DH78 | RNF224 | AF-P0DH78-F1-model_v3 | 95;112 | 0.49;1.0 |
| Q7L0R7 | RNF44 | AF-Q7L0R7-F1-model_v3 | 351;357;405;409;412 | 0.72;0.67;0.91;0.8;0.65 |
| Q9Y252 | RNF6 | AF-Q9Y252-F1-model_v3 | 80;608;630;644 | 0.35;0.75;0.44;0.62 |
| Q8IW03 | SIAH3 | AF-Q8IW03-F1-model_v3 | 23;25;45;173;213 | 0.97;0.7;0.91;0.07;0.5 |
| O15524 | SOCS1 | AF-O15524-F1-model_v3 | 118 | 0.18 |
| P40337 | VHL | AF-P40337-F1-model_v3 | 159;171;196 | 0.53;0.67;0.41 |

[a] Searched in human proteome, excluding sequences <150 aa, with the predicted number of TMH >2, or annotated as membrane-bound.
