## Supplementary figures and images for "Lysine-deficient proteome can be regulated through non-canonical ubiquitination and ubiquitin-independent proteasomal degradation"

### SFig1.pdf

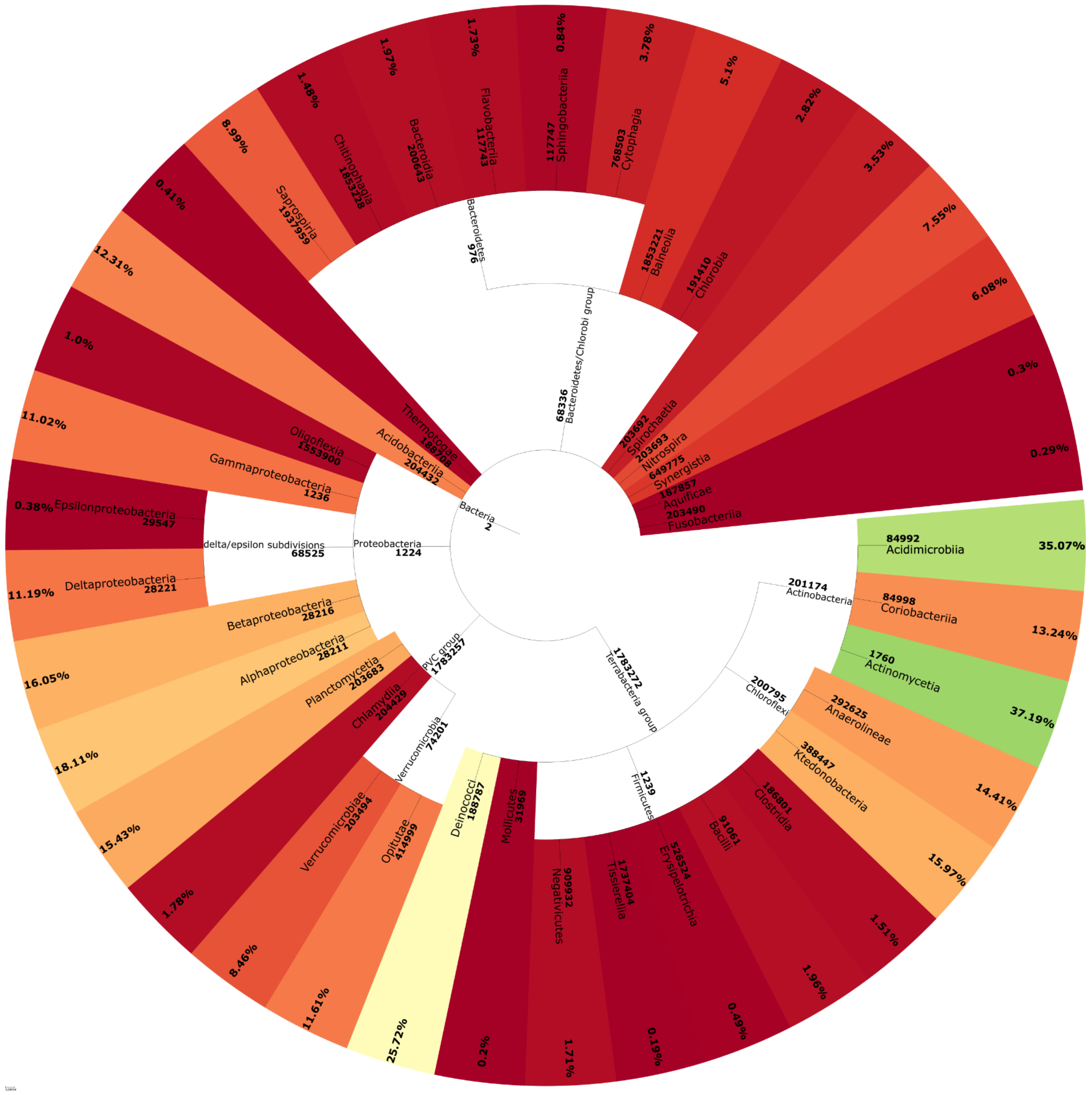

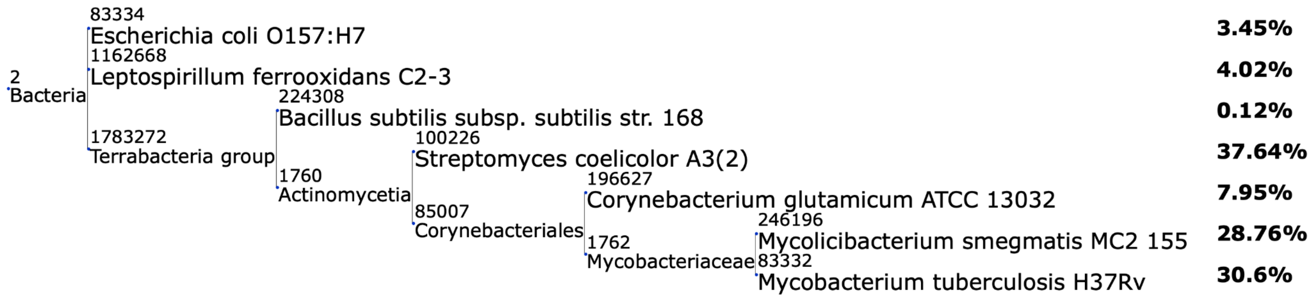

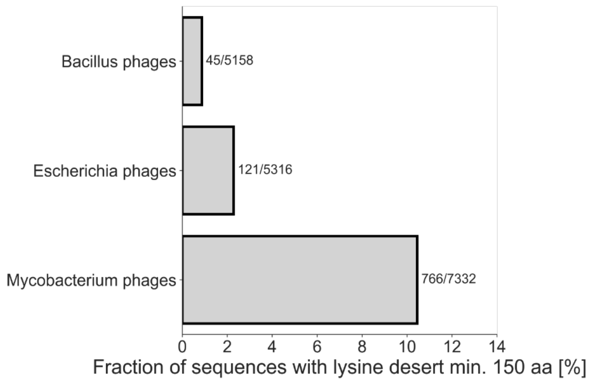

### SFig2.pdf

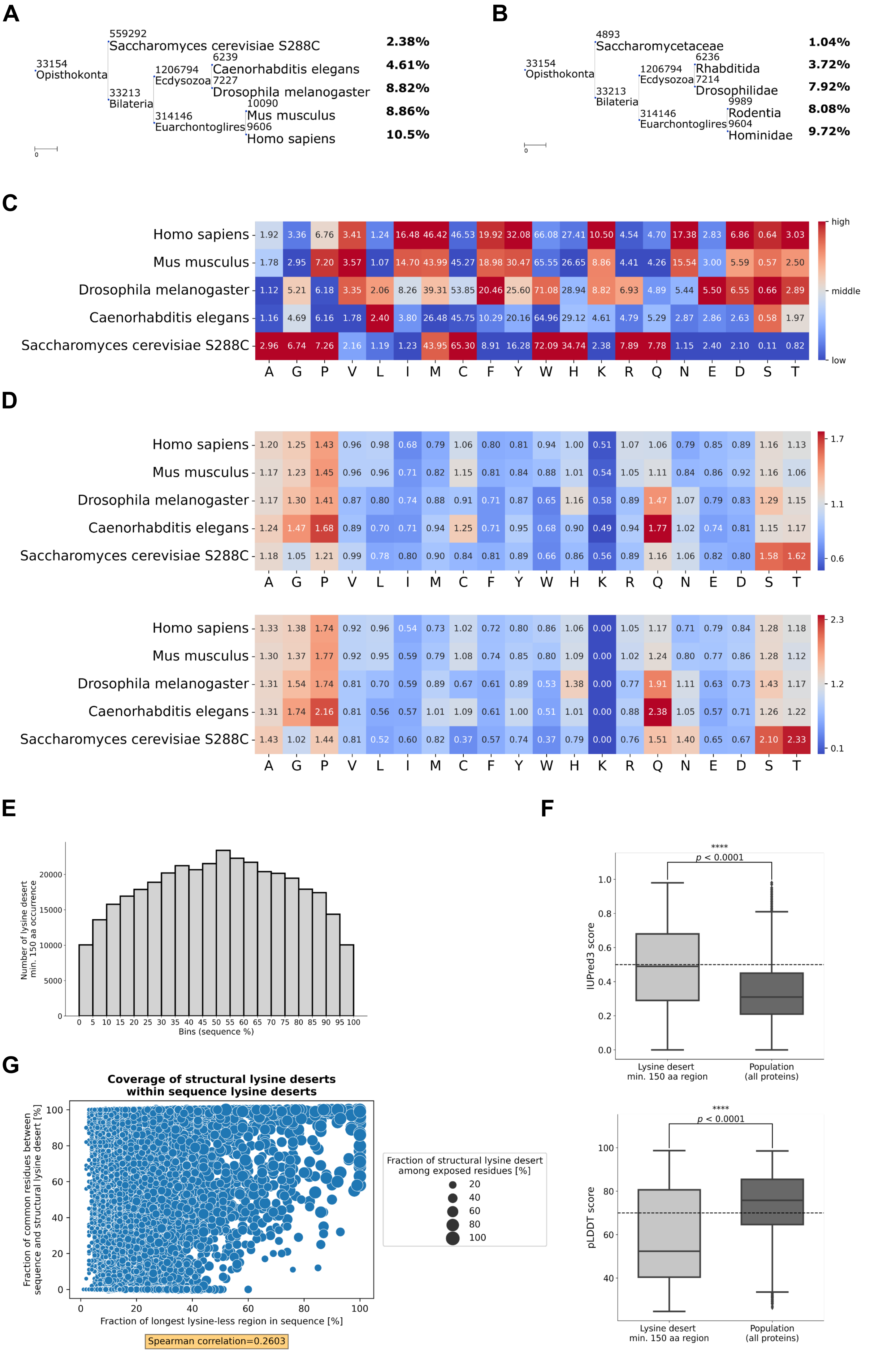

### SFig3.pdf

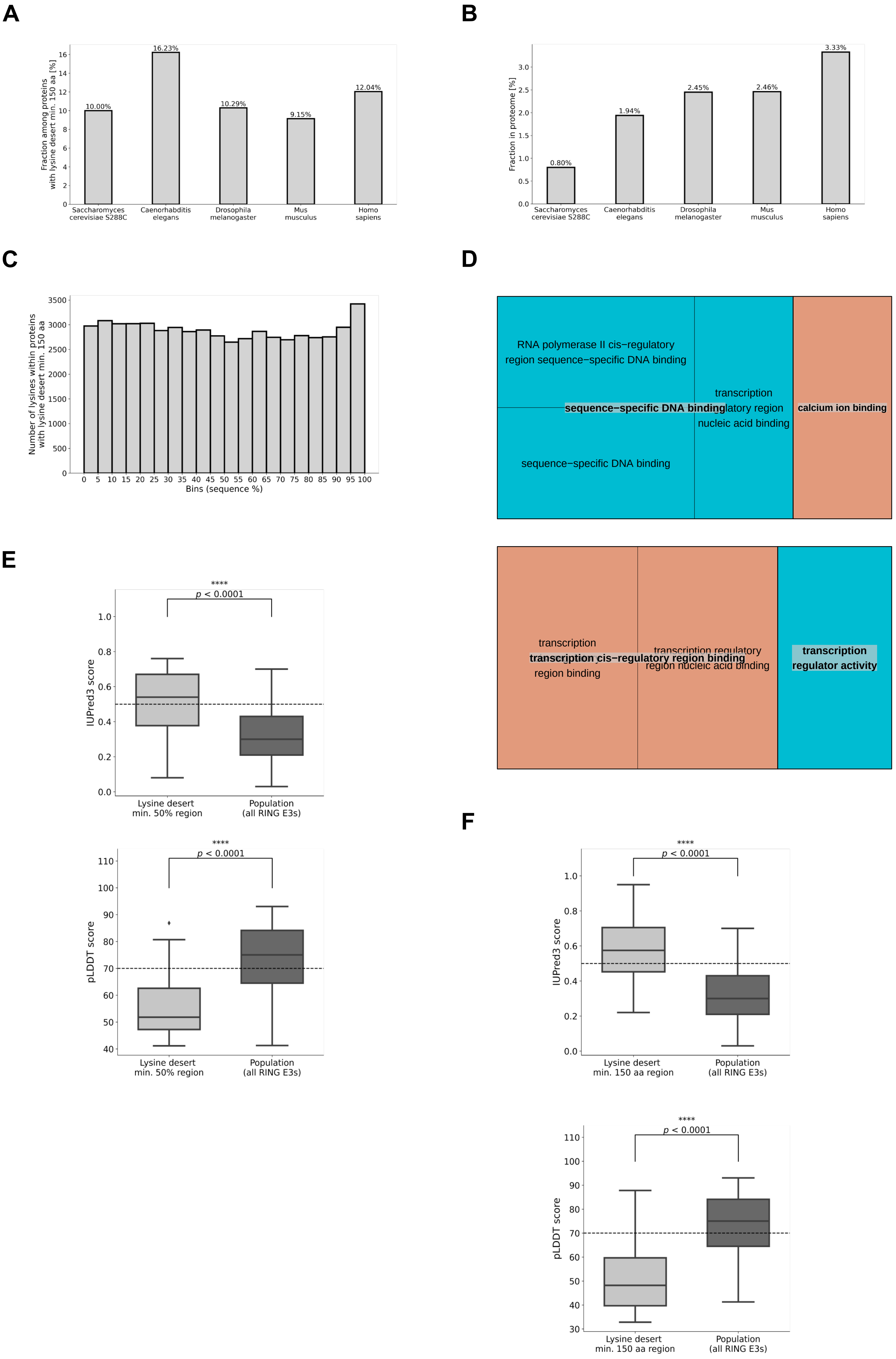

### SFig4.pdf

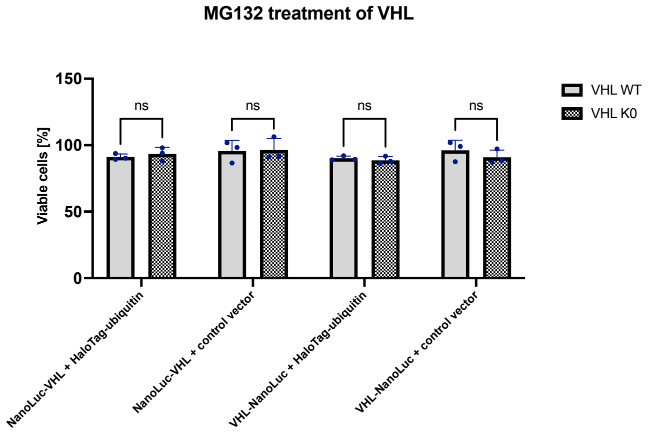

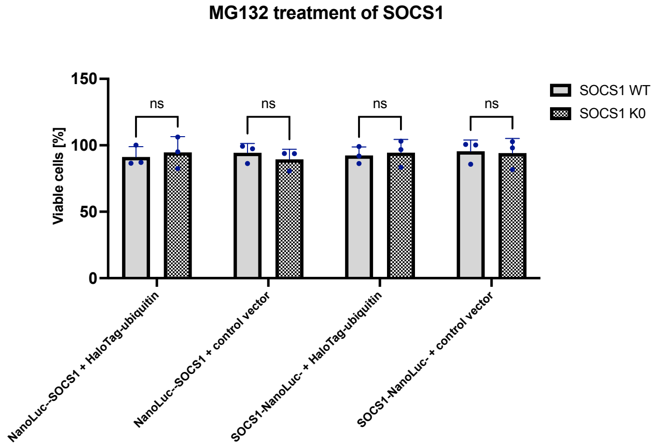

### SFig5.pdf

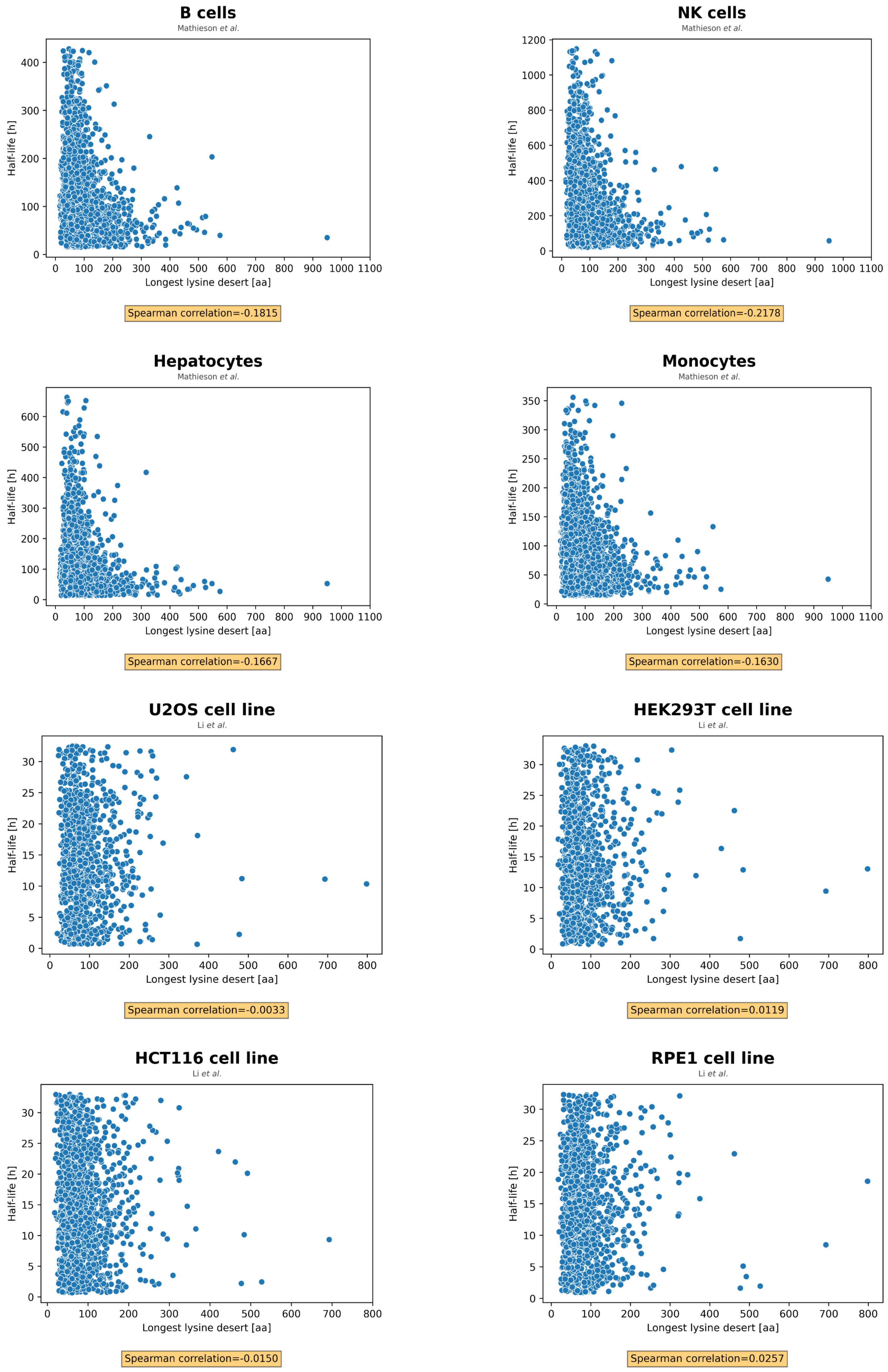
